# Interdependent RNA structural motifs at the 3ʹ-terminus of the West Nile virus genome regulate viral growth

**DOI:** 10.64898/2026.03.11.711107

**Authors:** Lucille H. Tsao, Doug E. Brackney, Anna Marie Pyle

**Affiliations:** Department of Chemistry, Yale University, New Haven, CT, 06511, USA; Department of Entomology, Connecticut Agricultural Experimental Station, State of Connecticut, New Haven, CT, 06511, USA; Department of Molecular, Cellular, and Developmental Biology, Yale University, New Haven, CT, 06511, USA; Howard Hughes Medical Institute, Chevy Chase, MD, 20815, USA

**Author notes:** Address correspondence to Anna Marie Pyle.

## Abstract

The RNA genome of West Nile Virus (WNV) folds into an elaborate series of RNA structural elements that are crucial for viral function. Among these elements, four pseudoknots (PKs) at the viral 3’-terminus, designated as SLII, SLIV, DBI, and DBII, are among the most crucial players in the overall flaviviral lifecycle. While many studies have focused on exploring the behavior of individual PKs, we investigated the collective role of all four PKs in viral growth and small flaviviral RNA (sfRNA) formation. Through mutational analyses and infectious models, we establish that the four PKs are interdependent and work synergistically to aid in the folding and compaction of the WNV 3’-terminal region. A striking hierarchy is observed in PK contributions to global folding and sfRNA formation, whereby SLIV plays the largest role, followed by DBI, DBII, and SLII. We also discover highly conserved RNA tertiary motifs within the PK assembly that are shared across flaviviruses, suggesting a new type of druggable target that may be of value in the search for pan-flaviviral therapeutics.

**GRAPHICAL ABSTRACT:** 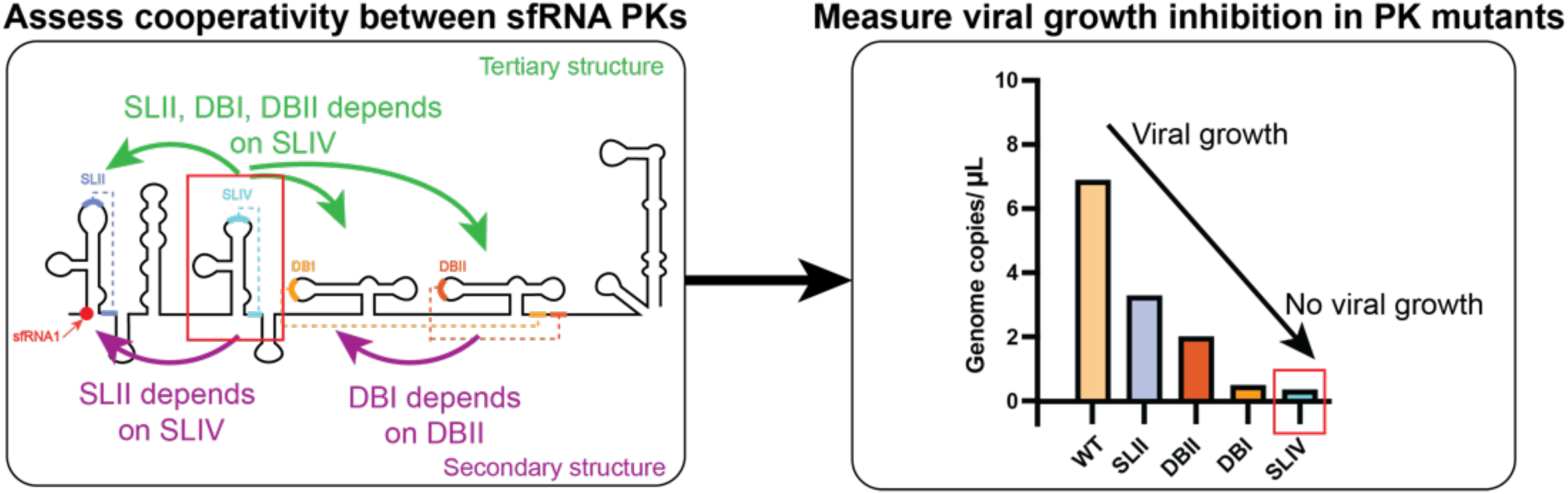

## INTRODUCTION

Flaviviruses pose one of the most significant challenges to human health, with approximately 400 million annual cases of infection worldwide. Flaviviruses (*Orthoflaviviridae*) include the widespread dengue (DENV) and Zika (ZIKV) viruses, as well as West Nile virus (WNV), which can lead to fatal neurological damage (1,2). Despite the human and economic toll of these viruses, the development of WNV or pan-*flaviviral* therapeutics remains particularly challenging due to the complexity of their genomes. One of the major obstacles to vaccine development is that deficient flavivirus genomes in live-attenuated vaccines can accumulate adaptive mutations and revert to the wild-type sequence (3–6). However, new vaccine strategies may be enabled by the fact that flaviviral genomes contain complex RNA structures that are critical to viral fitness (6–9). Modulating these structures, rather than sequence, has distinct therapeutic advantages (6).

Like other flaviviruses, WNV has a long (∼11 kb) positive-sense (+) RNA genome that contains complex secondary and tertiary RNA elements that facilitate innate immune evasion, host adaptation, and the viral life cycle. The WNV open reading frame (ORF), for instance, contains numerous riboregulatory RNA structural elements that can modulate viral growth (10). In addition, both the 5ʹ and 3ʹ untranslated regions (UTRs) of flaviviral genomes are comprised of conserved structural motifs that are crucial for regulating the viral life cycle and pathogenicity (11,12). Of particular importance, all flaviviruses produce a highly structured, non-coding subgenomic flaviviral RNA (sfRNA) that is derived from the 3ʹ UTR. These sfRNAs are formed through the partial degradation of the viral genomic RNA by the host 5ʹ-3ʹ exoribonuclease XRN1. Resistance to XRN1-mediated degradation results in the accumulation of sfRNAs, which impedes innate immunity and facilitates viral replication (13–20). That said, the precise mechanism of sfRNA action is not fully understood.

The structural organization of sfRNAs generally consists of four conserved pseudoknots (PKs) that have been identified in almost all types of flaviviral RNAs and have diverse functions in several viral RNA genomes (21–29). The four WNV sfRNA PKs are Stem-Loop (SL)-II, SLIV, Dumbbell (DB)-I, and DBII. The individual PKs and surrounding RNA regions are also referred to as xrRNAs, named for their ability to inhibit XRN1-mediated degradation (30). Together, the four PKs of WNV and other viruses create a highly compact and structured sfRNA (13). The largest WNV sfRNA is sfRNA1 (525 nt), which contains all four PKs. Smaller sfRNA products such as sfRNA2 and sfRNA3, which consist of 3 and 2 PKs, respectively, can also exist (11). Through pioneering crystallographic work, the structures of individual PKs have been well characterized and there is a good understanding of the complex relationship between the sfRNA and XRN1 (31,32). The functional importance of specific PKs has been actively investigated, establishing roles in protein recruitment or switching between translation and replication. Nevertheless, much of our understanding of sfRNA PKs has come from studies where PKs have been taken out of the full-length sfRNA context. While individual PKs can still form, we do not know the structural basis for how all PKs fold within an intact sfRNA and whether they coordinate with each other to mediate its formation and function.

Studies on RNA folding and compaction have established the vital role of magnesium ions (Mg^2+^) in defining RNA structure (33–35). Specific biophysical and biochemical work on sfRNAs has clearly demonstrated the importance of Mg^2+^ in stabilizing the long-range RNA-RNA interactions of the SLII PK so that it can mechanically resist XRN1-degradation (36,37). Nevertheless, PK dependence on Mg^2+^ is not fully understood, in part because a comprehensive analysis of sfRNA folding has not been undertaken.

Here we employ chemical probing methods such as Selective 2ʹ-Hydroxyl Acylation analyzed by Primer Extension and Mutational Profiling (SHAPE-MaP) and Terbium-seq (Tb-seq) as a function of Mg^2+^ to monitor the formation and stability of the four PKs in their natural, full-length context. Through mutational analysis, we evaluate the contribution of individual PK elements to intact sfRNA folding and virus infection, establishing a clear hierarchy in PK contribution to sfRNA folding. These studies establish that PKs are not isolated units, but rather they coordinate with each other and synergistically aid in the compaction of the sfRNA and the genomic 3ʹ-terminus. Our holistic assessment of sfRNA folding also revealed novel tertiary motifs that represent promising pan-flaviviral RNA drug targets. This integrated approach provides a blueprint for dissecting sfRNA function and mechanism, determining the crucial role of PK-PK interdependencies, and identifying potential therapeutic modules in all sfRNAs.

## MATERIAL AND METHODS

### Construct Design and Preparation

The wild-type (WT) West Nile virus (WNV) subgenomic-flaviviral RNA (sfRNA), 3ʹ terminus, and mutant sfRNA plasmids were derived from the WNV NY99 (AF404756) strain. Inserts containing the WT sfRNA or 3ʹ terminus sequence were obtained from Integrated DNA Technologies (IDT). Mutant WNV sfRNA constructs were generated by inverting the 5ʹ or 3ʹ end of the sequences involved in pseudoknot base pairing. Specifically, mutant SLII contains the following sequence changes: WT 5ʹ-A_55_CUCAAC-3ʹ sequence was replaced with 5ʹ-U_55_GAGUUG-3ʹ; mutant SLIV replaces 5ʹ-U_213_GC-3ʹ with 5ʹ-A_213_CA-3ʹ; mutant DBI replaces 5ʹ-A_403_ACACCA-3ʹ with 5ʹ-U_403_UGUGGU-3ʹ; and mutant DBII replaces 5ʹ-G_353_CUGU-3ʹ with 5ʹ-C_353_GACA-3ʹ. A list of all mutant constructs and their orientations can be found in Supplementary Figure S7. Mutant inserts were obtained through Quintara Biosciences and ThermoFischer. All constructs were cloned into pcDNA3.1 via Gibson Assembly Master Mix (NEB, cat #E2611S). Generated plasmids were then transformed into DH5-alpha competent cells and grown at 25°C overnight. Resulting plasmids were confirmed through Quintara Whole Plasmid Sequencing. Gibson Assembly primers can be found in Supplementary Table S1.

The WT full-length (FL) WNV plasmid contains the infectious cDNA clone of WNV NY99 (AF404756) (10). The FL WNV mutants contain mutations that disrupt secondary and tertiary structure. Specifically, mutant SLII contains the following sequence changes: WT 5ʹ-A_10557_CUCAAC-3ʹ sequence was replaced with 5ʹ-U_10557_GAGUUG-3ʹ; mutant SLIV replaces 5ʹ-U_10715_GC-3ʹ with 5ʹ-A_10715_CA-3ʹ; mutant DBI replaces 5ʹ-A_10905_ACACCA-3ʹ with 5ʹ-U_10905_UGUGGU-3ʹ; and mutant DBII replaces 5ʹ-G_10855_CUGU-3ʹ with 5ʹ-C_10855_GACA-3ʹ; mutant cyclization sequence replaces 5ʹ-A_10921_GCAUAUUGACA-3ʹ with 5ʹ-A_10921_GCUAUAAGACA-3ʹ; mutant Tb-DBI shortens 5ʹ-A_10800_CUAGAG-3ʹ with 5ʹ-A_10800_C-3ʹ; mutant Tb-DBII shortens 5ʹ-G_10873_ACUAG-3ʹ with 5ʹ-A_10873_C-3ʹ; mutant single strand control replaces 5ʹ-A_10744_AACCAAC-3ʹ with 5ʹ-A_10744_CCAAACA-3ʹ; and using synonymous mutations, mutant Domain I replaces 5ʹ-A_10311_GCCCGAGAACACGGUUAGUCCGAC-3ʹ with 5ʹ-C_10311_GCACGAGAACAAGGAUACUCACUA-3ʹ. A list of all mutant constructs and their orientations can be found in Supplementary Figure S6. Inserts were obtained from Quintara Biosciences. After double restriction enzyme digest, mutant inserts were ligated into a backbone derived from the WT FL WNV plasmid using T4 DNA ligase (NEB, cat #M0202S). Generated plasmids were transformed into HB101 competent cells (Promega, cat #L201B) and grown at 25°C overnight. Selected plasmids were verified through Quintara Whole Plasmid Sequencing. A list of all mutant constructs can be found in Supplementary Figure S7.

### WNV sfRNA and 3ʹ terminus *in vitro* transcription, semi-native purification, folding, and SHAPE-modification

WT and mutant sfRNA and the 3ʹ-terminus plasmids containing sequences as described above were linearized with Xba1 (WT, SLII and DBI mutants, 3ʹ-Terminus) (NEB, cat #R0145S) and EcoRI (SLIV and DBII mutants) (NEB, cat #R3101S). The linearized plasmid served as a template for *in vitro* transcription in conditions previously described (38). Briefly, the template was transcribed by T7 RNA polymerase P266L variant (39), followed by RQ1 DNase treatment (Promega, cat #M6101) and 5 ul of 0.5 M EDTA (pH 8) to chelate Mg^2+^. Transcribed RNA was further concentrated on 50 kDa Amicon Ultrafiltration columns and purified by size-exclusion on S200 increase GL 10/300 column with a 24 mL bed volume (Cytiva, cat # 28990944) at room temperature. The column was equilibrated with filtration buffer (40 mM 4-(2-hydroxyethyl)-1-piperazineethanesulfonic acid potassium salt [K-HEPES] pH 7.2, 150 mM KCl, 0.1 mM EDTA). RNA from the peak fraction was then diluted with filtration buffer to 100 ng/ul and folded by incubating with various Mg^2+^ concentrations (0, 0.5, 1, 3, 5, 10, and 15 mM Mg) at 37°C for 30 min. To probe the folded RNAs, a final concentration of 200 mM 2-methylnicotinic acid imidazolide (NAI, synthesized as described in (40)) or Dimethyl sulfoxide (DMSO) was added and incubated at 37°C for 10 min. The reaction was quenched and precipitated with ethanol and resuspended in 1 X ME buffer (10 mM morpholinepropanesulfonic acid [MOPS] pH 6 and 1 mM EDTA).

### *In vitro* SHAPE-MaP RT, Library Preparation, Quantification, and Sequencing

*In vitro*-purified probed RNAs (1 μg) were reverse transcribed (RT) with 200 U SuperScript II (SSII) (ThermoFisher, cat #18064014) under SHAPE-MaP SSII buffer conditions (50 mM Tris-HCl [pH 8.3], 75 mM KCl, 10 mM dithiothreitol [DTT], 6 mM MnCl_2_, and 0.5 mM deoxynucleoside triphosphate [dNTP] mix) using gene specific primers (Supplementary Table S1). RT reactions were done at 42°C for 3 hours. cDNAs were then purified with AmpureXP beads (Beckman Coulter, cat #A63881) with a 1.8:1 bead to sample ratio. Purified cDNAs were then diluted to 0.2 ng/ul for library preparation. Sequencing libraries were generated using the Nextera XT DNA library preparation kit (Illumina, cat #FC-131-1096) following manufacturer’s protocols. Libraries were quantified using Qubit (Life Technologies) and TapeStation (Agilent) and sequenced on an Illumina NextSeq 2000 platform.

### SHAPE-MaP Data Analysis and Structure Prediction

All SHAPE-MaP libraries were analyzed using ShapeMapper2.1.5 (41). The sequences were aligned to the WNV sfRNA WT/mutant or 3ʹ-Terminus sequence derived from NY99 (AF404756). Default alignment parameters and QC metrics were used. All libraries passed the three ShapeMapper quality control metrics (Supplementary Table S2).

The secondary structures for all constructs were predicted by Superfold (42). Default parameters were used. A list of PK and single stranded region constraints included in SuperFold predictions are found in Supplementary Table S3. The constraints used were based on previous *in silico* predictions and functional data (11,12,43). The structure outputs from SuperFold were visualized in StructureEditor (44) and RNAcanvas (45). Reactivities and predicted structures were compared to *in cellulo* WNV structure obtained previously (10) to assess whether *in vitro* structure was useful for further study.

To track changes in PK formation upon structural mutation, normalized SHAPE reactivities for the pseudoknotted nucleotides found in each PK in the mutants and WT were averaged. Then the difference between the mutant and WT average reactivities (ΔSHAPE) at each Mg^2+^ titration point was calculated (Figure 3; Supplementary Figure S3A-D). For each heat map, a symmetric color gradient was set relative to the largest absolute difference, with ΔSHAPE=0 indicating no difference between mutant and WT average reactivities. Positive changes in green correspond to higher reactivity (i.e. more modified), while negative changes in purple correspond to lower reactivity (i.e. less modified).

### *In Vitro* Tb-seq, RT, and Library Preparation

*In vitro* Tb-seq was performed as described (46). Briefly, WT and mutant sfRNAs were transcribed and purified as described above. Purified RNA was then diluted to 50 ng/ul and folded at various Mg^2+^ concentrations (0, 1, 3, 5 mM Mg) at 37°C for 30 min. To probe the folded RNAs, a final concentration of 0.5 mM Tb^3+^ (Sigma-Aldrich, cat #212903-5g) was added and incubated at room temperature for 10 min. For a denaturing control, formamide was added to 50% to folded RNA before adding 0.5 mM Tb^3+^. All reactions were quenched with 3 ul of 50 mM EDTA pH 8. Probed RNAs were cleaned with Zymo RNA Clean and Concentrator (Zymo Research, cat #R1017) following manufacturer’s protocol and eluted with water.

*In vitro*-purified probed RNAs (1 μg) were mixed with 2 pmol of gene-specific primers (Supplementary Table S2). Primers were annealed to RNAs by heating to 90°C for 1 min and 30°C for 2 min. The RNAs were then reverse transcribed (RT) with MarathonRT (RNAConnect), 2X MarathonRT buffer, (100 mM Tris-HCl pH 8.3, 400 mM KCl, 4 mM MgCl2, 10 mM DTT, 40% glycerol) and 0.5 mM dNTP (NEB, cat #N0447S). RT reactions were incubated at 42°C for 30 minutes. RNA was then digested with 0.5 ul of RNase T1 (ThermoFisher, cat #EN0541), H (NEB, cat #M0297S), and A (NEB, cat #T3018L) and incubated at 37°C for 30 min. cDNAs were then purified with AmpureXP beads (Beckman Coulter, cat #A63881) with a 1.2:1 bead to sample ratio. A final concentration of 0.5 uM 3ʹ adaptor was then ligated to purified cDNAs (Supplementary Table S1), 0.5 mM ATP, 10 U T4 RNA ligase, 1X T4 RNA ligase buffer (NEB, cat #M0204S), and 25% PEG 8000. The reaction was incubated at 25°C for 16 hours, followed by enzyme deactivation at 65°C for 15 min. Ligated products were then purified with AmpureXP beads with a 1.2:1 bead to sample ratio. Purified products were Q5 PCR (NEB, cat #M0491S) amplified with Illumina TruSeq and indexed reverse primers (NEB Next Multiplex Oligos, cat # E7335S, E7500S, E7710S, E7730S). Libraries were quantified using Qubit (Life Technologies) and bioanalyzer (Agilent) or TapeStation (Agilent), diluted, pooled, and sequenced on a NextSeq 2000 platform (Illumina).

### Tb-seq Data Analysis

All scripts to analyze Tb-seq libraries can be found in a GitHub repository: https://github.com/pylelab/Tb-seq. The analysis and quality checks were performed following the pipeline previously described (46). Briefly, using RTEventsCounter, sequences were aligned to the WT or mutant WNV sfRNA sequence. Using RStudio, the intensity of Tb cleavage sites was normalized against the denaturing control. To confidently identify Tb cleavage sites, z scores for each nucleotide normalized intensity were first calculated. Then, the following general guidelines were used to identify Tb cleavage sites:

1. Any nucleotides with a z score less than 1 were excluded as Tb cleavage sites.
2. Any nucleotides found on stems were excluded as Tb cleavage sites, except for nucleotides at the base of stems.
3. Tb cleavage sites must consist of a cluster of 2 or more nucleotides.

To track changes to Tb-sites upon structure disruption, the normalized Tb-seq reactivities for the nucleotides found in each Tb-site were averaged. Reactivities were visualized on a heat map, where the color gradient was set relative to the WT Tb-seq average reactivities. The standard error of the mean was also calculated for each Tb-seq average reactivity. Mutant average reactivities that were closer to the WT average reactivity indicated that a Tb-site was preserved and were colored green (tertiary motif present, allowing for more Tb-induced cleavage), while low averages indicated a Tb-site was abolished and were colored red (no tertiary motif present, no Tb-induced cleavage) (Supplementary Figure S5). The secondary structures and Tb cleavage sites for all constructs were visualized in StructureEditor (44) and RNAcanvas (45).

### Tb-site Sequence Conservation Analysis

We compiled 3ʹ UTR sequences extracted from the NCBI genome database of the most common mosquito-borne flaviviruses (Supplementary Table S4). We also included several mosquito-borne flaviviruses from the Japanese encephalitis serocomplex to better survey sequence conservation of flaviviruses closely related to WNV. The sequences were then aligned against the WNV 3ʹ UTR sequence using the UniPro UGENE Align Tool (Clustal Omega) with default settings. The sequences of WNV Tb-sites were compared to the other aligned sequences. If all nucleotides from each Tb-site sequence were conserved, the site was considered 100% conserved. More than one nucleotide differences compared to the WNV Tb-site sequence were excluded and considered not conserved.

### WNV FL *in vitro* transcription and capping

The WT FL WNV and mutant FL WNV plasmids were linearized using Xba1 (NEB, cat #R0145S), followed by ethanol precipitation and resuspended in water. The linearized plasmids served as templates for *in vitro* transcription in conditions as described above for WNV sfRNA transcription. Afterwards, RNAs were purified using RNeasy kit following manufacturer’s protocols (Qiagen, cat #74104). RNAs were then capped using the Vaccinia capping Enzyme (NEB, cat #M2080S) and mRNA Cap 2ʹ-O-Methyltransferase (NEB, cat #M0366S) according to manufacturer’s protocols. The capping protocol added a type-1 cap with an inverted 7-methylguanosine and 2ʹ-OMe on the first nucleotide. Capped RNAs were then purified again using the RNeasy kit (Qiagen cat #74104) and eluted in 1X ME buffer.

### Cell Culture

Vero cells (American Type Culture Collection [ATCC], CCL-81) were cultured according to ATCC guidelines. Briefly, cells were cultured with Dulbecco’s Modified Eagle Medium (DMEM) supplemented with 10% heat-inactivated fetal bovine serum (HI-FBS), nonessential amino acids (NEAA), L-glutamine, and sodium bicarbonate. Vero cells were incubated at 37°C/5% CO_2_. Experiments using live WNV were all performed in the Connecticut Agriculture Experimental Station’s BSL3 facility.

### q-RT-PCR Viral Growth Assay adapted from (10,47)

A quantitative RT-PCR assay was used to quantify viral growth. The assay was adapted from (47) and previously described (10). Briefly, a 2.5 kb RNA standard (Genome coordinates: 1031-3431). PCR template was created from the plasmid that contains the full-length infectious cDNA clone of WNV. This standard that contains the E gene was then *in vitro* transcribed following the same protocol as described above. The standard was then purified using RNeasy columns (NEB, cat #74104) following manufacturer protocols and eluted in 1XME. After the RNA was quantified, it was aliquoted in 10-fold serial dilutions.

Vero cells were seeded in 12-well plates at 2.5E5 cells per well and left to grow overnight. Then, 500 ng of capped genomic RNAs were transfected into Vero cells using TransIT-mRNA Transfection Kit (Mirus, cat # MIR 2225). Four hours post-transfection, wells were washed with 1X Phosphate Buffered Saline (PBS) and supplemented with 10% FBS-DMEM. Supernatant samples were collected at 4 to 6 days post infection (dpi). Supernatant samples were then cleaned with RNase A for 30 minutes at 37°C followed by Proteinase K treatment for 1 hours at 37°C. RNA was isolated using the MagMAX Prime Viral/Pathogen NA Isolation Kit (ThermoFischer, cat # A58145) and a KingFischer 711 Automated Extraction and Purification System (Thermo).

To determine the viral copy number of collected supernatant samples, we used a 6-carboxy-fluorescein (FAM)-labeled TaqMan probe (ThermoFisher) that targets the E gene and the Luna Universal Probe One-Step RT-qPCR kit (NEB, cat #E3006S). Viral RNA genome copy per microliter of sample across 2 replicates was calculated using a linear regression derived from the standard curve. A one-way ANOVA with multiple comparisons was performed across 10 groups. All statistics were performed using tests available in GraphPad Prism. Primers can be found in Supplementary Table 2.

## RESULTS

### Secondary structure of sfRNA from WNV successfully recapitulated *in vitro*

According to previous in cell SHAPE-MaP studies on the full length WNV genome (10), the 3ʹ terminus of the WNV genome contains three domains that adopt a compact RNA architecture (Figure 1A). To assess whether these domains adopt a similar fold outside the context of the full-length genome, SHAPE-MaP was performed on an *in vitro* transcribed RNA construct containing the 3ʹ terminal regions studied in isolation (Figure 1A, Domains I-III). We then performed a constrained secondary structure prediction using Superfold (Supplementary Table 3) (42). The resulting secondary structure prediction of the isolated 3ʹ terminus demonstrated a striking similarity to the same region found in Vero and C6/36 cells infected with WNV (Supplementary Figure S1A) (10). Notably, the *in vitro* secondary structure implies that the sfRNA region (Figure 1A, also known as Domain III) is likely to be the most structured section of the 3ʹ-terminus, consistent with previous observations *in cellulo* (10). To determine if the isolated sfRNA region does indeed adopt the same structure in isolation as it does within the 3ʹ terminus and in the context of the full length genome in infected cells, we *in vitro* transcribed a construct containing only the sfRNA in isolation. We then performed SHAPE-MaP and compared the calculated secondary structure of the isolated sfRNA to the full length 3ʹ-terminal construct. Analysis of the isolated sfRNA secondary structure revealed a comparable number of protected nucleotides (<0.4 SHAPE reactivity) as that observed for the sfRNA region when it is embedded within the larger construct (67% vs. 66%) (Figure 1B). Additionally, we observed a strong correlation in nucleotide reactivities between sfRNA in isolation with those observed for the sfRNA region *in cellulo* from both C6/36 (Figure 1C, r = 0.81) and Vero (Figure 1D, r = 0.80). Taken together, these results indicate that the secondary structure of the sfRNA studied *in vitro* is nearly identical to the same region within the entire genomic RNA in infected cells, indicating it folds autonomously.

**Figure 1.**
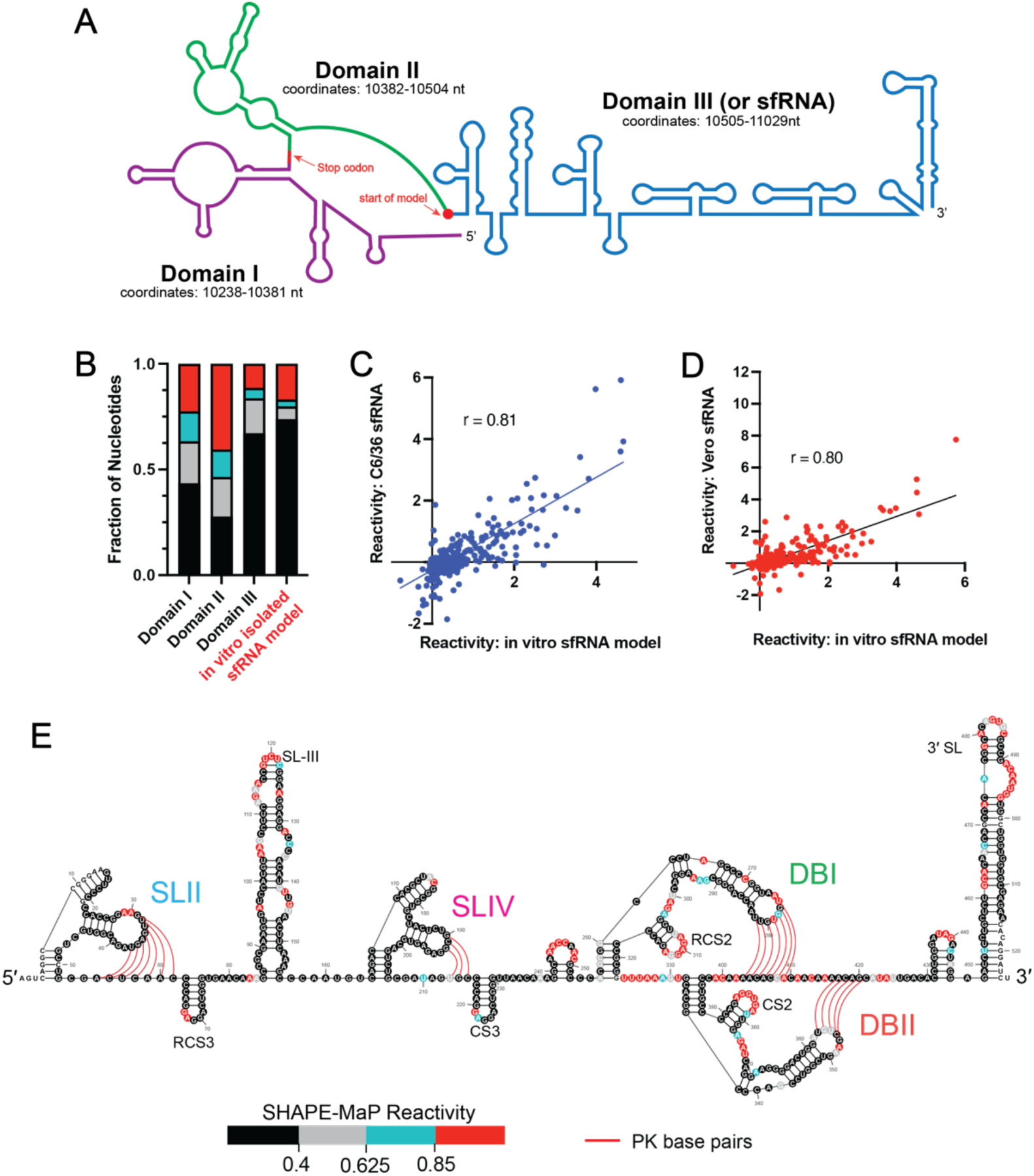
Compact sfRNA secondary structure is observed *in cellulo* and *in vitro*. (***A***) Schematic of 3’ viral terminus in WNV. The 3’ terminus is composed of three structural domains: Domain I (purple), Domain II (green), and Domain III (blue). Domain III is more commonly known as the sfRNA. The stop codon and the start of the isolated sfRNA construct are indicated with the red arrows. (***B***) Nucleotides from Domains I, II, and III in the *in vitro* 3’ terminus construct and from the isolated sfRNA construct were grouped by normalized reactivity. Each group is expressed as a fraction of the total number of nucleotides found in each Domain. (***C, D***) Comparison of normalized SHAPE reactivities between the *in vitro* sfRNA construct to the sfRNA region found in Vero (C) and C6/36 (D) cells. Lines represent linear regressions fit to the data. The Pearson’s correlation for each comparison is displayed. (***E***) Averaged normalized SHAPE reactivities from two replicates of the *in vitro* sfRNA construct folded at 1 mM Mg^2+^ mapped onto Superfold predicted secondary structure map of the sfRNA. Pseudoknot base pairs are represented with red lines. SHAPE reactivities are categorized based on the following thresholds: <0.40 (black), 0.40≤ to <0.625 (gray), 0.625≤ to <0.85 (blue), ≥0.85 (red).

We compared the experimentally determined secondary structure model of the sfRNA in isolation (Supplementary Figure S1B) with previously published *in silico* and genetically predicted models of the WNV sfRNA secondary structure (Figure 1E) (11,12,43). This analysis revealed that all previously reported structural motifs, such as the four pseudoknots, conserved hairpins, and the 3ʹ SL are successfully recapitulated within our experimentally determined sfRNA structure and the in-cell structure (Figure 1E) (11,43). The correspondence of this data enabled us to establish a biologically relevant construct containing isolated sfRNA that could be used for mutational and functional analyses at the biochemical level.

### sfRNA pseudoknots form in a hierarchical manner

Consistent with previous work, our experimentally determined sfRNA secondary structure is composed of 4 pseudoknots (PKs) known as Stem-Loops (SL)-II and IV, and Dumb-Bells (DB)-I and II, each of which is presumed to play a role in sfRNA folding (Figure 1E). We set out to quantitatively determine the relative stability of the four pseudoknots simultaneously by using SHAPE-MaP to monitor the formation of sfRNA base pairings as a function of magnesium ion concentration ([Mg^2+^]) (Figure 2A; Supplementary Figure S2) (48). In these experiments, monovalent cation was also provided to support the basic secondary structure of duplex stems (150 mM Potassium Chloride, KCl), but tertiary interactions (such as pseudoknot base-pairings) are not expected to form in the absence of Mg^2+^ (33,49,50). Our global analysis reveals important overall trends in sfRNA folding where the individual sfRNA pseudoknots have different Mg^2+^ requirements for folding. In the baseline secondary structure obtained at 0 mM Mg^2+^, 42-67% of the PK nucleotides were susceptible to SHAPE modification (≥0.85 SHAPE reactivity, red) (Figure 2A(i)), suggesting that none of the PK domains are completely folded in the absence of Mg^2+^. Upon addition of 0.5 mM Mg^2+^, the SLII pseudoknot nucleotides became completely protected (<0.40 SHAPE reactivity, black), indicating stable SLII formation (Figure 2A(ii)). By contrast, the other three PKs are incompletely folded at 0.5 mM Mg^2+^, as demonstrated by the presence of both partially protected (0.4 – 0.625 SHAPE reactivity, gray) and unprotected nucleotides (red). To achieve a uniformly folded conformation, SLIV and DBII require 1 mM Mg^2+^ (Figure 2A(iii)), while DBI requires more than 1 mM Mg^2+^ (Supplementary Figure S2).

**Figure 2.**
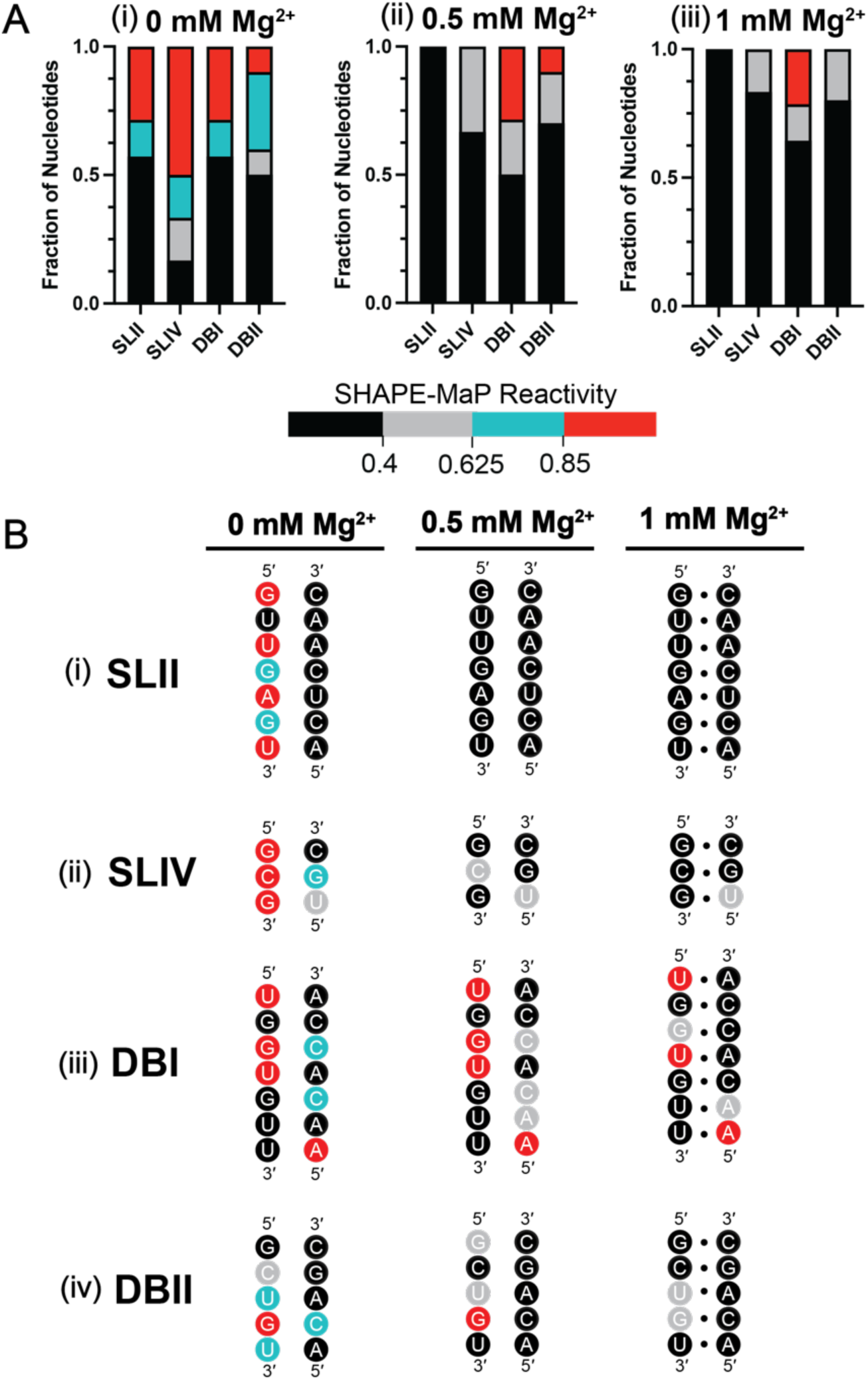
Mg^2+^-dependent folding of sfRNA PKs. (***A***) Normalized reactivities for each pseudoknotted nucleotide found in SLII, SLIV, DBI, and DBII PKs were averaged from two replicates at 0, 0.5, and 1 mM Mg^2+^. Nucleotide reactivities in each PK were then grouped by reactivity level (i.e. amount of SHAPE modification) based on the following thresholds: <0.40 (black, low levels of modification), 0.40≤ to <0.625 (gray, partial to low levels of modification), 0.625≤ to <0.85 (blue, partial to high levels of modification), ≥0.85 (red, high levels of modification). For each PK, the pseudoknotted nucleotides were then expressed in fraction of total plots to display the distribution of SHAPE reactivities found in each PK. (***B***) Sequences of pseudoknotted nucleotides at 0, 0.5, and 1 mM Mg^2+^. Pseudoknotted nucleotides from the four PKs are colored based on average reactivity level across two replicates: low reactivity (black), low-medium reactivity (gray), high-medium reactivity (blue), high reactivity (red).

Analysis of nucleotide-level dependence of individual PK secondary structures at each Mg^2+^ concentration provides a more detailed picture (Figure 2B). For example, we observed that at 0 mM Mg^2+^, the only structures formed in any of the PK structures are local duplex stems. Although these stems are protected from SHAPE modification, nucleotides involved in long-range PK base pairings are susceptible to SHAPE modification, indicating that the PK pairings have not formed under this condition (Figure 2B(i)-(iv), 0 mM Mg^2+^). Upon increasing the Mg^2+^ concentration to 0.5 mM, SLII appears to be the only PK that forms all its base-pairs (Figure 2B(i), 0.5 mM Mg^2+^). SLIV is the next most stable PK, as it contains nucleotides that are partially protected from SHAPE modification under this condition (0.4 - 0.625 SHAPE reactivity, gray) (Figure 2B(ii), 0.5 mM Mg^2+^). At this Mg^2+^ concentration, both DBI and DBII contain nucleotides that are susceptible to modification (Figure 2B(iii), (iv), 0.5 mM Mg^2+^, red). At 1 Mg^2+^, PK base pairing increases for both SLIV and DBII, indicating complete PK formation (Figure 2B(ii), (iv), 1 mM Mg^2+^). By contrast, DBI requires much higher concentrations of Mg^2+^ (at least 5 mM) for all PK base pairs to form (Figure 2B(iii), 1 mM Mg^2+^; Supplementary Figure S2). Taken together, these results indicate that individual PKs have different levels of stability. SLII is the most stable PK, forming at the lowest [Mg^2+^]. The next most stable PK is SLIV, followed by DBII, and finally, DBI is the least stable structural element.

### Pseudoknots in the sfRNA are interdependent and jointly contribute to its overall structure

Given the hierarchical nature of PK stability, we sought to determine whether PKs function autonomously or if they are interdependent. To investigate this, we created four mutant sfRNA transcripts in which each individual PK is unfolded by the incorporation of mutations that are expected to maximally disrupt base pairing of a single, individual PK (Supplementary Figure S6). SHAPE-MaP was then performed over a range of [Mg^2+^] to gauge whether the disruption of one PK influences the Mg^2+^ dependency of the other PKs. To quantitate the resulting effects, we calculated the difference between the mutant and WT SHAPE reactivities (ΔSHAPE) at each [Mg^2+^] (Figure 3). For example, a ΔSHAPE=0 indicates no difference between mutant and WT reactivities at a given position. Positive changes (shown in green, Figure 3) correspond to higher reactivity (i.e. more modified) in the mutant, while negative changes correspond to lower reactivity (purple, i.e. less modified). To evaluate how individual sequence mutants altered the global sfRNA architecture, we supplemented our analysis with secondary structure predictions for each mutant transcript at 0.5 (Supplementary Figure S3) and 1 mM Mg^2+^ (Supplementary Figure S4).

**Figure 3.**
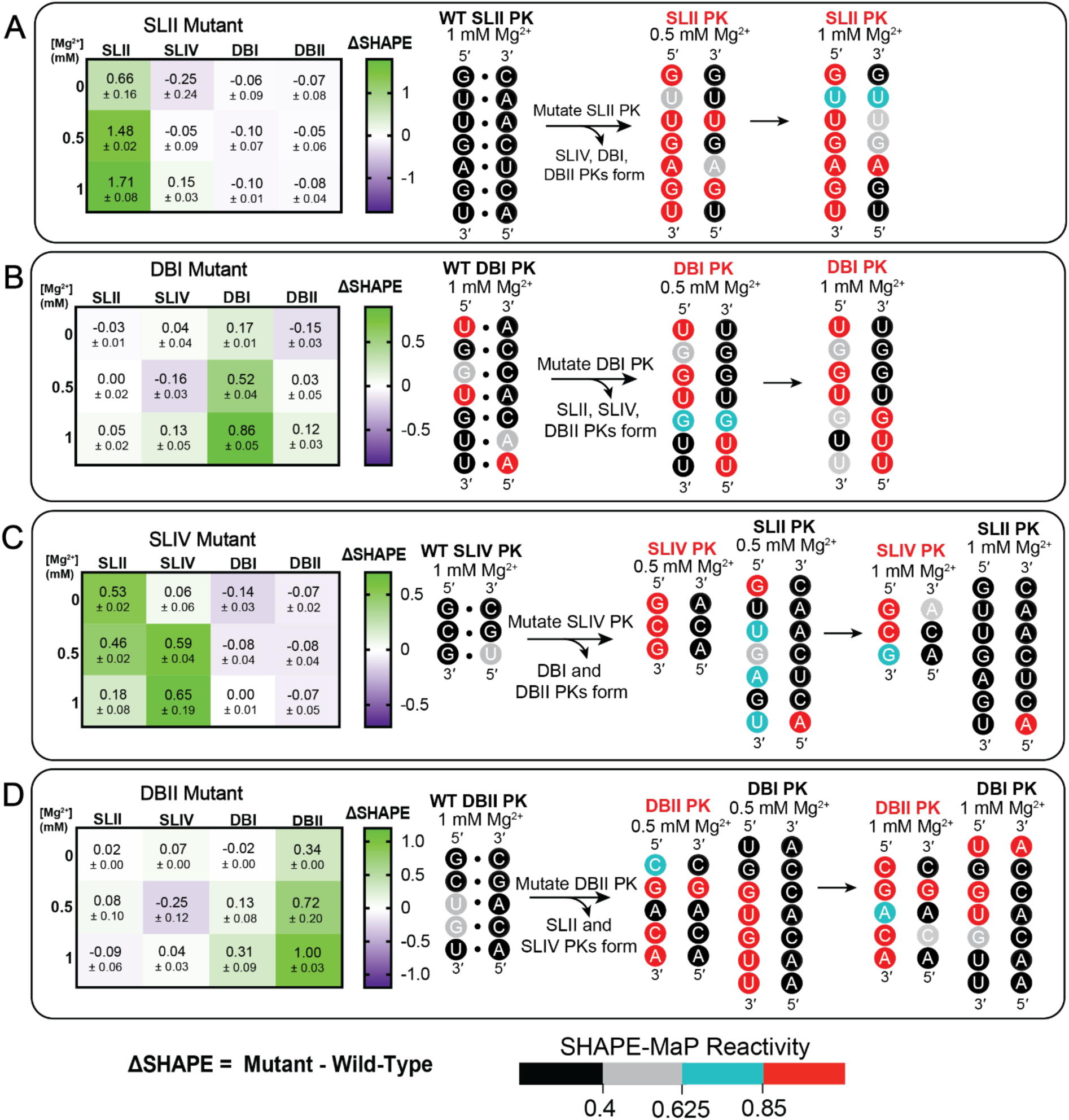
Mutational analysis of the sfRNA reveals pseudoknot interdependence. (***A-D***) The differential average normalized SHAPE-MaP reactivity of pseudoknotted nucleotides (ΔSHAPE = mutant – WT) for each sfRNA mutant (SLII, SLIV, DBI, DBII) was calculated (two replicates). The symmetric color gradient for each heat map was determined by the largest absolute differential reactivity. Where ΔSHAPE = 0, there is no difference between the mutant and WT reactivities. Positive changes (green) correspond to higher reactivity (more modified), while negative changes (purple) correspond to lower reactivity (less modified). Errors represent the standard error of the mean. Next to each heat map are schematics representing nucleotide changes to specific PKs when mutations are introduced at 0.5 and 1 mM Mg^2+^. Nucleotides are shaded or circled based on the following reactivity thresholds: (<0.40 (black, low levels of modification), 0.40≤ to <0.625 (gray, partial to low levels of modification), 0.625≤ to <0.85 (blue, partial to high levels of modification), ≥0.85 (red, high levels of modification).

Taken together, these analyses demonstrated that mutations in the SLII PK base pairs had no effect on the stability of the other three PKs and led to negligible changes in global sfRNA folding (Figure 3A, left heat map; Supplementary Figure S3A, S4A). The mutated SLII PK was unable to form PK base pairs at 0.5 and 1 mM Mg^2+^, compared with the WT SLII PK structure at the same concentrations (Figure 3A, right schematic; Figure 2B(i)). Likewise, mutation of the DBI PK base pairings did not lead to changes to the stability of the other constituent PKs (Figure 3B; Figure 2B(iii); Supplementary Figure S3C; S4C). By contrast, mutations of the SLIV PK pairings induced the most dramatic effects on the neighboring PKs (Figure 3C). Unzipping of SLIV causes SLII to become destabilized, unable to properly base pair at 0.5 mM Mg^2+^ (Figure 3C, right schematic; Supplementary Figure S3B). While in the WT context the SLII PK was originally observed to stabilize at 0.5 mM Mg^2+^ (Figure 2B(i)), it now required 1 mM Mg^2+^ to reach a more stable state (Figure 3C, right schematic; Supplementary Figure S4B). Additionally, disruption to SLIV did not appear to affect DBI and DBII formation (Figure 3C, left heat map; Supplementary Figure S3B; S4B). Lastly, DBII mutation destabilized the formation of DBI but not SLII and SLIV (Figure 3D, left heat map). Detailed analysis of nucleotide reactivities show that DBI base pairing is compromised at 0.5 and 1 mM Mg^2+^ (Figure 3D, right schematic; Supplementary Figure S3D, S4D) when compared to the base pairing seen in the WT (Figure 2B(iii)). Overall, a sequential mutational analysis indicates that the PKs do not fold independently of each other. Instead, they appear to work synergistically to facilitate overall sfRNA folding.

### Higher-ordered motifs near sfRNA pseudoknots drive sfRNA tertiary folding

SHAPE analysis of sfRNA secondary structural features suggests that the PKs do not function as isolated units, but rather they behave as interdependent modules within a larger tertiary structure. To monitor global sfRNA tertiary structure more directly, we employed Terbium-induced cleavage (Tb-seq), which reveals positions of sharp, tight backbone turns that are characteristic elements of compact RNA tertiary structural motifs and RNA-protein interactions (46). Tb-seq was performed on *in vitro* transcribed WT sfRNA at 5 mM Mg^2+^, which is a condition in which all four PKs are expected to be completely formed (Supplementary Figure S2). Under this condition, twelve regions of strong Terbium cleavage (Tb-sites) are observed (Figure 4A, green nucleotides, 1-12; Supplementary Figure S5A), a majority of which are located near DBI and DBII PKs (Figure 4A, sites 3 – 11, nt 290 to 420). Two sites were also found near SLII and SLIV PKs (Figure 4A, site 1 at nt 76-79 and site 2 at nt 112-113), and one was observed at the 3ʹ Stem-Loop (3ʹ SL) (Figure 4A, site 12 at nt 453-455).

**Figure 4.**
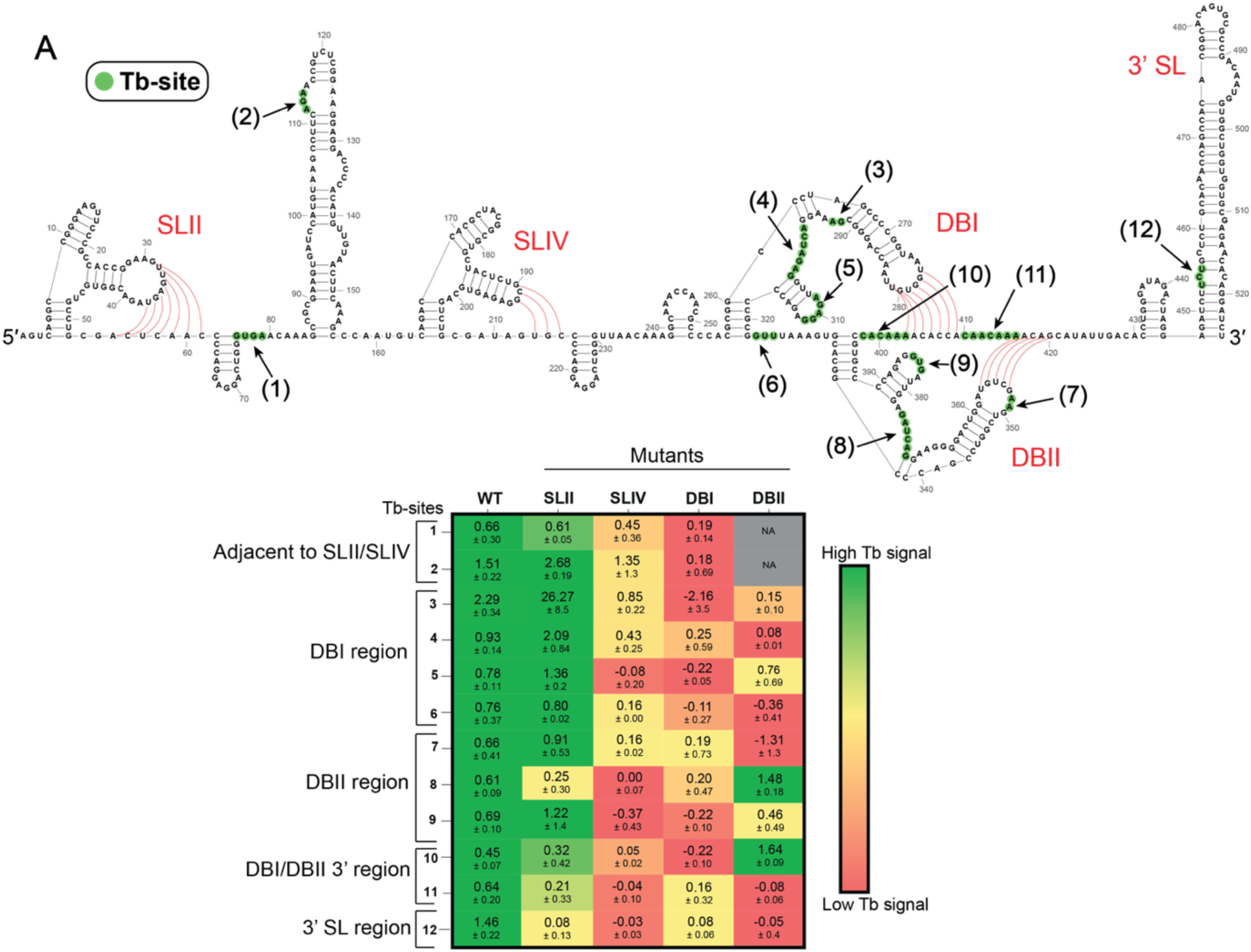
Pseudoknot couplings contribute to sfRNA tertiary folding. (***A***) Tb-seq signals were averaged across two replicates. Nucleotides corresponding to a Tb-site are highlighted with green circles on the secondary structure map of the WT sfRNA. Each Tb-site is labelled (1) to (12) in black and indicated with black arrows. Each PK is labelled in red, and pseudoknot base pairs are shown with red lines. (***B***) Tb-seq was performed on the four mutant sfRNA constructs. To compare the mutant results to the WT, the Tb-seq signals for each nucleotide in each of the 12 Tb-sites were averaged across two replicates to obtain an average Tb-signal for the WT and mutants. A green to red gradient based on the average Tb-signals was then applied on the mutants with the WT serving as the reference. Tb-signals in the mutant that were equal or greater to the signal found in WT are colored green, indicating the presence of a Tb-site. Low Tb-seq signals, indicating no Tb-site, are colored red. Errors represent the standard error of the means.

Having identified distinct Tb-sites in the WT sfRNA, we then performed Tb-seq on the four mutant sfRNA constructs described above, monitoring whether individual PK mutants preserved (green), destabilized (yellow), or abolished (red) each of the twelve Tb-sites (Figure 4B; Supplementary Figure S5B-E). This analysis suggests that the SLII mutant does not strongly influence overall tertiary fold of the sfRNA, as 10 Tb-sites are preserved in the SLII mutant (Figure 4B, sites 1-7, 9-11; Supplementary Figure S5B). By contrast, mutation of the other PKs induced major effects on overall tertiary organization. For example, mutation of SLIV caused sites near the SLII/SLIV and DBI regions to diminish (Figure 4B, sites 1-7) and sites near the DBII region were completely abolished (Figure 4B, sites 8-12; Supplementary Figure S5C). Similarly, the DBI mutant abolished all Tb-sites near SLIV (Figure 4, sites 1-2) and DBII (Figure 4, sites 7-10). Strikingly, mutation of DBII abolished Tb-sites in the SLII, SLIV, and DBI regions (Figure 4, sites 1-2, 5-7, 9, and 11), but preserved Tb-sites 8 and 10. Taken together, these results indicate that the SLIV, DBI, and DBII PKs contribute to formation of the overall sfRNA tertiary structure. Furthermore, these results reveal new couplings between SLIV, DBI, and DBII at the tertiary level, aligning with our understanding that PK interdependencies promote sfRNA folding.

### WNV sfRNA PK tertiary motifs are conserved across flaviviruses

The preceding Tb-seq analysis suggests that the four PKs within the WNV sfRNA are components of a larger tertiary structural unit. Given that sfRNA secondary structures and pseudoknots are broadly conserved across flaviviruses (51), we sought to investigate conservation of tertiary organization across mosquito-borne flaviviruses by examining the sequence conservation of Tb-sites. To this end, we performed a sequence alignment between 3ʹ UTR sequences of flaviviruses from different serocomplexes and WNV WT 3ʹ UTR (Figure 5B; Supplementary Table 4) (52). The results indicate that the WT WNV sfRNA Tb-site sequences are conserved across flaviviral 3ʹ UTRs (Figure 5A; Supplementary Figure S6). Specifically, 4 Tb-sites located near DBI and DBII (sites 4, 5, 7, and 8, Figure 5A) are strongly conserved across all flaviviruses analyzed. Within the DBI region, site 4 is 100% conserved in all members of the Japanese encephalitis virus (JEV) serocomplex (Figure 5C). We also found a single nucleotide difference in the four dengue virus (DENV) serotypes. Site 5 was 100% conserved in both the JEV and DENV serocomplexes. Neither site was conserved in Zika virus (ZIKV) or yellow fever virus (YFV) (Figure 5C). Similarly, in the DBII region, site 7 was 100% conserved in both the JEV and DENV serocomplexes but not in ZIKV or YFV (Figure 5D). Notably, site 8 is the most conserved Tb-site, as it was 100% conserved in JEV and DENV serocomplexes and ZIKV, and partially conserved in YFV (Figure 5D; Supplementary Figure S6). These data suggest that the motif located at site 8 is an interesting candidate as a pan-flaviviral therapeutic target.

**Figure 5.**
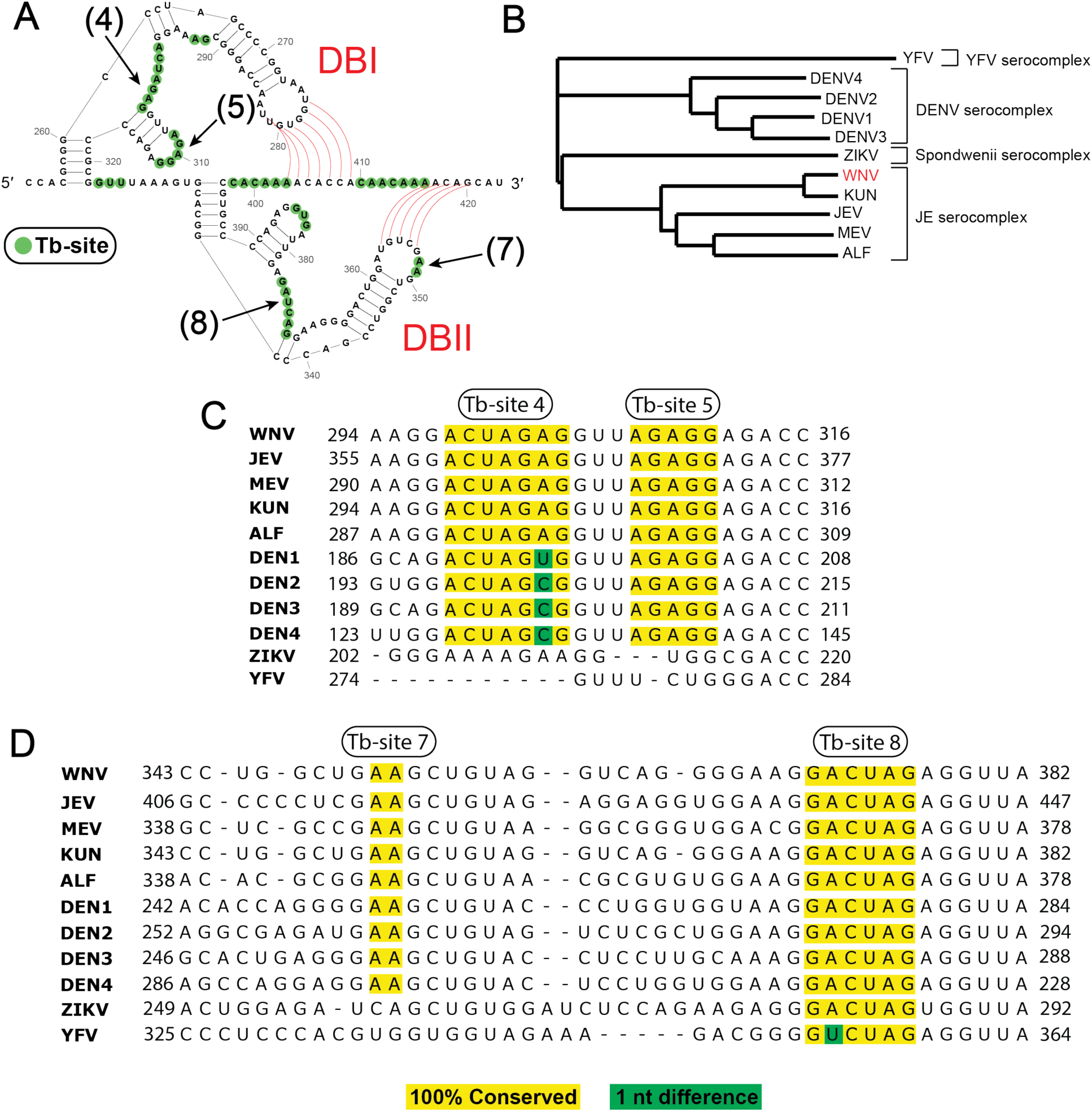
sfRNA Tb-site sequences are conserved across flaviviruses. (***A***) Schematic showing Tb-sites found in WT sfRNA DBI and DBII PKs. Nucleotides found in each Tb-site are highlighted with a green circle. The specific sites used for sequence alignment are labeled in black and indicated with black arrows (sites 4, 5, 7, and 8). DBI and DBII PKs are indicated in red, and pseudoknot base pairs are shown with red lines. (***B***) Phylogenetic tree of flaviviruses used in conservation analysis. Serocomplexes are labeled with brackets. Tree generated using Unipro UGENE Build Tree tool. (***C, D***) The sequence conservation of DBI Tb-sites 4 and 5 (B) and DBII Tb-sites 7 and 8 (C) was performed by first obtaining several flaviviral 3’ UTR sequences from NCBI. A multiple sequence alignment was performed using Unipro UGENE Align Tool (Clustal Omega). WNV served as the reference sequence. Tb-sites that were found to be 100% conserved in a flavivirus 3’ UTR are highlighted in yellow. Single nucleotide differences are highlighted in green.

### sfRNA folding paradigm facilitates WNV viral growth integrity

Given the apparent synergy between the four PKs and the conservation of their surrounding tertiary structures, we evaluated their function in the context of a viral infection. To this end, we designed 8 full length WNV constructs that contain mutations in key structural motifs examined in the biochemical analyses. We then monitored how each of these mutant viruses infect and grow in cells (Supplementary Figure S7). Four of the viruses contain mutations in PK arm regions, where they are expected to maximally disrupt PK base pairing (Supplementary Figure S7). An additional variant contains synonymous mutations within the 3ʹ-terminal ORF region known as Domain I (nt 10238 – 10381), which are expected to disrupt secondary structure without affecting translation (10,53). Mutations were incorporated into the two highly conserved Tb-sites, Tb-DBI (based on Tb-site 4, Figure 5) and Tb-DBII (based on Tb-site 8, Figure 5) by deleting portions of the Tb-site sequence, thereby interfering with local tertiary structure (Supplementary Figure S7). We also designed the positive SS control mutant (nt 10744 – 10751) by scrambling the sequence of a highly SHAPE modified single-stranded region of the sfRNA (Figure 1; Supplementary Figure S7). We also included a negative control, Cyc_Def, which is a variant known to prevent genome cyclization and viral growth (10). All mutants and WT full length WNV were then *in vitro* transcribed, Type 1 capped and transfected into Vero cells. To measure changes in viral growth, we quantified viral RNA copy number in supernatant samples using an established q-RT-PCR protocol (10,47). Comparison of the mutants to the WT and SS control showed that all mutant constructs led to a significant reduction in viral copy number (Figure 6A), highlighting their importance for viral function. The data indicate that disruption of SLII does not inhibit viral growth as effectively as the other mutants. Based on the level of viral growth inhibition for the various mutants, we were able to determine a hierarchy for the contribution of each PK to sfRNA function: SLIV > DBI > DBII > SLII (Figure 6B), which aligns well with the biochemical analyses. Additionally, both Tb-seq mutants led to comparable growth inhibition as the negative control Cyc_Def mutant, suggesting that these two mutants are highly detrimental to viral function and could serve as pan-flaviviral drug targets. Overall, these results indicate that disruption of sfRNA 2D and 3D structural motifs severely inhibits viral growth and that SLII is not a major contributor to functional secondary and tertiary sfRNA formation.

**Figure 6.**
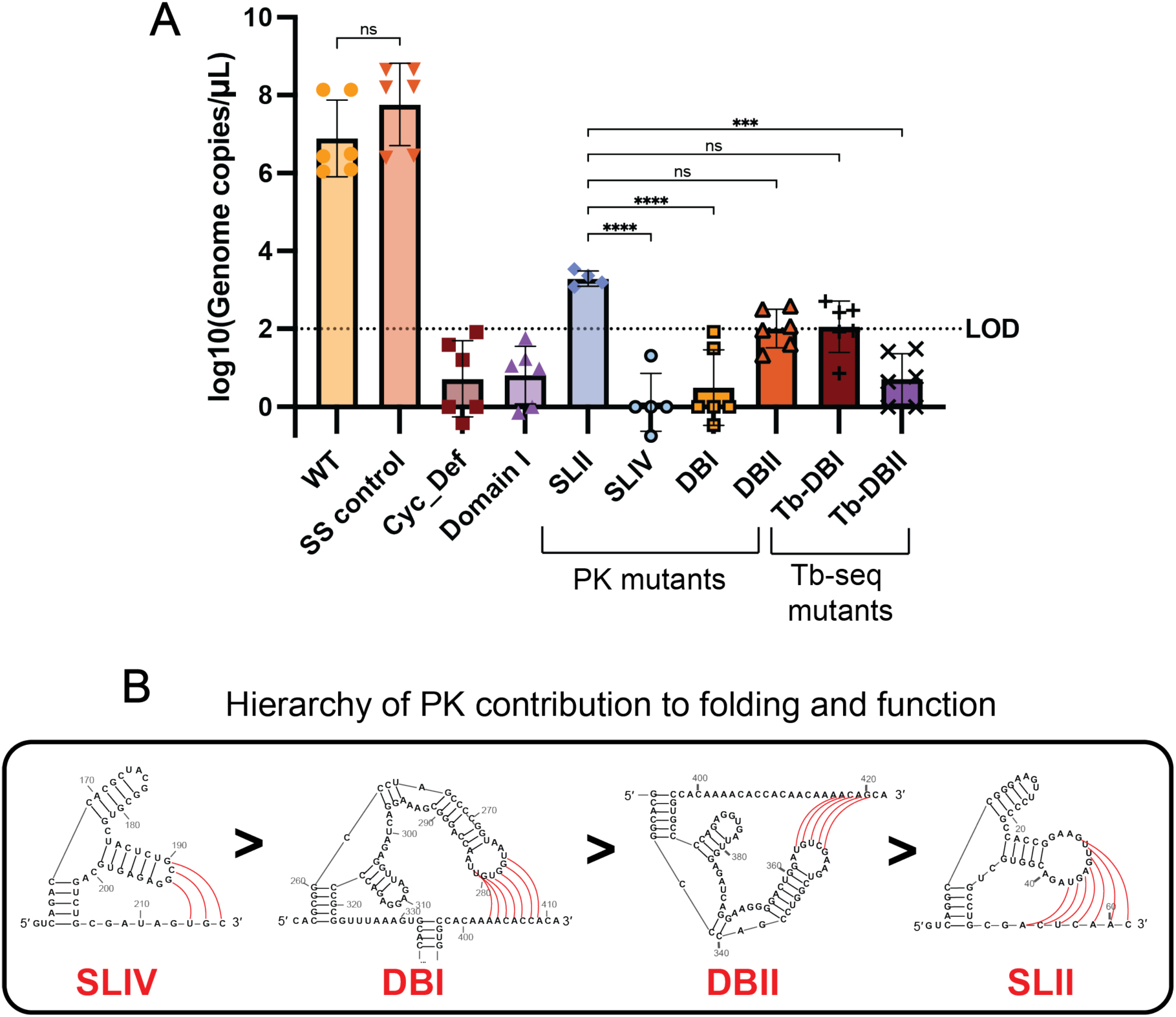
Viral growth defects resulting from RNA motif mutants. (***A***) Viral growth was measured by quantifying viral genomes in cell supernatant with q-RT-PCR in Vero cells at 5 dpi. Each condition had 6 independent technical replicates. The bars represent the average viral growth and error bars represent the standard deviation. A one-way ANOVA with multiple comparisons was performed (ns: not significant, *P < 0.05, **P < 0.01, ***P < 0.001, ****P < 0.0001). (***B***) A proposed PK hierarchy for how each PK contributes to sfRNA function was determined based on levels of viral growth inhibition. The hierarchy shows that SLIV > DBI > DBII > SLII. Secondary structures of each PK are displayed. Pseudoknot base pairs are shown in red.

## DISCUSSION

While it is generally accepted that structural features at the 3ʹ-end of flaviviral genomes are crucial to viral function (13–16,18,20), many structural and functional details remain unknown. In particular, it has been unclear whether the constituent PKs interact with each other to influence the folded state of the genomic terminus and the corresponding sfRNAs derived from it. The current model of sfRNA function (15,31,32,54) suggests that sfRNAs adopt a set of tightly folded structures that resist degradation by XRN1. Prevailing models suggest that an sfRNA remains unfolded until SLII, the upstream PK, forms a three-way junction that enables the rest of the sfRNA to subsequently fold and compact (15). In certain ways, our results align with that model, as we observe that SLII is the most autonomously stable PK (Figure 2). However, our mechanistic analysis of overall sfRNA folding reveals that the story is more complicated. By systematically evaluating the influence of individual sfRNA components on overall architecture, folding paradigm, and function of the WNV 3ʹ-terminus, we observe hierarchical and interactive formation of PKs, each of which makes important contributions to viral function. The multiple orthogonal approaches employed in this study show that all four PKs contribute to 3ʹ-end stability and sfRNA architecture with the following hierarchy of functional and structural importance: 1) SLIV, 2) DBI, 3) DBII, and 4) SLII. These findings establish that individual PKs are not autonomous units; rather they work synergistically to facilitate optimal folding of a functional architectural unit at the downstream terminus of flaviviral genomes.

In terms of sfRNA folding hierarchy, our results suggest a modification of the previously proposed paradigm in which the rigid SLII element plays a central role in viral immune evasion, serving as the first line of defense against XRN1-mediated degradation (11,13,54,55). The data presented in our study suggests that SLII plays a more peripheral role in overall sfRNA compaction. Instead, we observe that SLIV behaves as a master regulator of global sfRNA folding, modulating the stability and formation of the other constituent PKs. SLIV formation may restrict the conformational landscape of sfRNA folding, thus providing the other three PKs with the proper geometry for folding into a functional, stable structure. Indeed, when present in the same transcript, SLIV influences the stability of SLII folding, resulting in a coupled process that promotes efficient sfRNA production and viral growth. These data suggest that SLIV may play a critical role in aiding WNV host adaptation between humans and mosquitoes, as reported in studies of DENV (3,4,56). We also observe striking interplay between DBI and DBII and that the disruption of one DB PK destabilizes the other. In DENV, DBII can regulate the switch between the linearized (translation) and circularized (replication) form of the genome (22,27). It is therefore plausible that the WNV DB PKs may act as similar molecular switches between translation and replication.

Despite decades of biochemical study on flaviviral PK folding, there is little functional data on the role of PKs and other 3ʹ-flaviviral genome motifs during the process of viral infection. To address this gap in understanding, we used the biochemical SHAPE-MaP and Tb-seq findings to guide the design of novel, infectious WNV strains, employing viral growth assays to directly assess how individual PKs and other 3ʹ-terminal motifs impact WNV function. The experimental difficulty of these experiments cannot be overstated: giant flaviviral genomes (11 kb) are exceptionally difficult to clone (57–59), and the full-length WT and mutant WNV viruses examined here were necessarily studied under stringent Biosafety Level 3 (BSL3) conditions (60). Despite the higher complexity of viral infection studies, the mutational results were even more striking than in the biochemical experiments, indicating that interdependence of the PKs is particularly important in the context of active infection. Disruption of all four sfRNA PKs reduced viral growth in cells (Figure 6), but as observed biochemically, certain PKs appear to play a more central role than others. The viral growth studies are consistent with a model in which SLIV is a primary regulator of 3ʹ-end stability, as SLIV mutations lead to the largest inhibition of viral growth. By contrast, SLII mutations displayed the smallest level of viral growth inhibition. Moderate effects were observed for the intricately coupled DBI and DBII. As in the biochemical studies, in-cell infection studies with full-length virus strains recapitulate the functional hierarchy of interdependent modules whereby SLIV > DBI > DBII > SLII.

In addition to the four PK mutants employed in the infection assays, we designed two additional mutants that probe elements of sfRNA tertiary structure. Guided by the Tb-seq experiments, we mutated the highly conserved sequences we discovered at sites of strong Tb^3+^ cleavage. These mutants severely inhibit viral growth, and given their sequence conservation across flaviviruses, our findings suggest they could serve as pan-flaviviral RNA targets (Figure 6). This study underscores the power of high-throughput Tb-seq experiments for identifying target therapeutic candidates, as it bypasses the need to selectively disrupt individual base-pairings and it flags features that are common to all flaviviral serotypes (5,6,61). The discovery of functional, pan-flaviviral RNA structural elements represents a crucial step forward in understanding and combating flaviviruses. Importantly, the same method would be applicable to any other mRNA or viral RNA of therapeutic relevance.

In summary, we established that modular 3D substructures at the 3ʹ-terminus of flaviviral genomes play a crucial role in viral infection and that these elements operate synergistically, forming larger, stable structures that contribute to sfRNA formation, viral replication and possibly other functions. To examine the biological relevance of flaviviral substructures, we employed mutational functional analysis with infectious models, thereby establishing a promising new set of possible therapeutic targets (Figure 5; Supplementary Figure S6). It is notable that sfRNA-defective live attenuated vaccines are under active development for flaviviruses such as WNV, DENV, and ZIKV (6,62). Our findings suggest ways to use structure-based design to improve such vaccine constructs. The apparent architectural complexity of the WNV 3ʹ-terminus suggests that, in addition to vaccines and ASO methodologies, small-molecule antiviral therapies may also be possible. Indeed, parallel efforts to target viral untranslated regions (UTRs) with small molecules are showing great promise. For example, small-molecule inhibitors for hepatitis C virus (HCV) that target structures in the 5ʹ UTR have been developed (8,9,18). Other teams have successfully employed antisense locked nucleic acids (LNAs) to disrupt specific RNA structural motifs and inhibit the growth of viruses such as WNV, DENV, SARS-CoV-2, and HCV (10,63–66). Our work adds to this foundation, establishing that knowledge of viral RNA tertiary structures, and not just their protein-coding sequences, can help meet the global health challenge posed by viral infections and possibly other diseases.

## ACKNOWLEDGEMENTS

We thank Dr. Nicholas C Huston, Dr. Shivali Patel, and Dr. Han Wan for thoughtful guidance and discussions. We would also like to thank all current Pyle lab members for their helpful comments. Special thanks to Michael Van Zandt and New England Discovery Partners (NEDP) for synthesizing and donating the NAI SHAPE-MaP reagent.

## AUTHOR CONTRIBUTIONS

Lucille H. Tsao: Conceptualization, Formal analysis, Methodology, Investigation, Validation, Writing—original draft. Doug E. Brackney: Methodology, Writing—review & editing. Anna Marie Pyle: Conceptualization, Methodology, Supervision, Writing—original draft,

## SUPPLEMENTARY DATA

Supplementary Data are available at NAR online.

## DATA AVAILABILITY

All data are available in the main text or the supplementary materials. SHAPE-MaP and Tb-seq data from this study have been deposited in GEO Accession Database PRJNA1427911.

## REFERENCES

1. Pierson, T.C., Diamond, M. S. (2020) The continued threat of emerging flaviviruses. Nature Microbiology, 5, 796–812.

2. Whitehorn, J., Yacoub, S. (2019) Global warming and arboviral infections Clinical Medicine, 19, 149–152.

3. Villordo, S.M., Filomatori, C.V., Sánchez-Vargas, I., Blair, C.D. and Gamarnik, A.V. (2015) Dengue Virus RNA Structure Specialization Facilitates Host Adaptation. PLOS Pathogens, 11, e1004604.

4. Filomatori, C.V., Carballeda, J.M., Villordo, S.M., Aguirre, S., Pallarés, H.M., Maestre, A.M., Sánchez-Vargas, I., Blair, C.D., Fabri, C., Morales, M.A. et al. (2017) Dengue virus genomic variation associated with mosquito adaptation defines the pattern of viral non-coding RNAs and fitness in human cells. PLOS Pathogens, 13, e1006265.

5. Araujo, S.C., Pereira, L.R., Alves, R.P.S., Andreata-Santos, R., Kanno, A.I., Ferreira, L.C.S. and Gonçalves, V.M. (2020) Anti-Flavivirus Vaccines: Review of the Present Situation and Perspectives of Subunit Vaccines Produced in Escherichia coli. Vaccines, 8, 492.

6. Doets, K. and Pijlman, G.P. (2024) Subgenomic flavivirus RNA as key target for live-attenuated vaccine development. Journal of Virology, 98.

7. Le Grice, S.F. (2015) Targeting the HIV RNA genome: high-hanging fruit only needs a longer ladder. 2015/03/05 ed.

8. Dibrov, S.M., Parsons, J., Carnevali, M., Zhou, S., Rynearson, K. D., Ding, K., Garcia Sega, E., Brunn, N. D., Boerneke, M. A., Castaldi, M. P., Hermann, T. (2014) Hepatitis C virus translation inhibitors targeting the internal ribosomal entry site. Journal of Medicinal Chemistry, 57, 1694–1707.

9. Hermann, T. (2016) Small molecules targeting viral RNA. Wiley Interdisciplinary Reviews: RNA, 7, 726–743.

10. Huston, N.C.T., L. H.; Brackney, D. E.; Pyle, A. M. (2024) The West Nile virus genome harbors essential riboregulatory elements with conserved and host-specific functional roles. Proceedings of the National Academy of Sciences, 121, 12.

11. Pijlman, G.P., Funk, A., Kondratieva, N., Leung, J., Torres, S., van der Aa, L., Liu, W. J., Palmenberg, A. C., Shi, P. Y., Hall, R. A., Khromykh, A. A. (2008) A highly structured, nuclease-resistant, noncoding RNA produced by flaviviruses is required for pathogenicity. Cell Host and Microbe, 4, 579–591.

12. MacFadden, A., O’Donoghue, Z., Silva, Pagc, Chapman, E. G., Olsthoorn, R. C., Sterken, M. G., Pijlman, G. P., Bredenbeek, P. J., Kieft, J. S. (2018) Mechanism and structural diversity of exoribonuclease-resistant RNA structures in flaviviral RNAs. Nature Communications, 9, 119.

13. Funk, A., Truong, K., Nagasaki, T., Torres, S., Floden, N., Balmori Melian, E., Edmonds, J., Dong, H., Shi, P. Y., Khromykh, A. A. (2010) RNA structures required for production of subgenomic flavivirus RNA. Journal of Virology, 84, 11407–11417.

14. Göertz, G.P., Fros, J.J., Miesen, P., Vogels, C.B.F., Van Der Bent, M.L., Geertsema, C., Koenraadt, C.J.M., Van Rij, R.P., Van Oers, M.M. and Pijlman, G.P. (2016) Noncoding Subgenomic Flavivirus RNA Is Processed by the Mosquito RNA Interference Machinery and Determines West Nile Virus Transmission by Culex pipiens Mosquitoes. Journal of Virology, 90, 10145–10159.

15. Slonchak, A., Khromykh, A. A. (2018) Subgenomic flaviviral RNAs: What do we know after the first decade of research. Antiviral Research, 159, 13–25.

16. Schuessler, A., Funk, A., Lazear, H. M., Cooper, D. A., Torres, S., Daffis, S., Jha, B. K., Kumagai, Y., Takeuchi, O., Hertzog, P., Silverman, R., Akira, S., Barton, D. J., Diamond, M. S., Khromykh, A. A. (2012) West Nile virus noncoding subgenomic RNA contributes to viral evasion of the type I interferon-mediated antiviral response. Journal of Virology, 86, 5708–5718.

17. Murira, A., Lamarre, A. (2016) Type-I interferon responses: from friend to foe in the battle against chronic viral infection. Frontiers in Immunology, 7, 609.

18. Vicenzi, E., Pagani, I., Ghezzi, S., Taylor, S. L., Rudd, T. R., Lima, M. A., Skidmore, M. A., Yates, E. A. (2018) Subverting the mechanisms of cell death: flavivirus manipulation of host cell responses to infection. Biochemical Society Transactions, 46, 609–617.

19. Pan, Y., Cai, W., Cheng, A., Wang, M., Yin, Z., Jia, R. (2022) Flaviviruses: innate immunity, inflammasome activation, inflammatory cell death, and cytokines. Frontiers in Immunology, 13, 829433.

20. Martin, M.F., Nisole, S. (2020) West nile virus restriction in mosquito and human cells: a virus under confinement. Vaccines, 8, 256.

21. Pallarés, H.M., Costa Navarro, G.S., Villordo, S.M., Merwaiss, F., De Borba, L., Gonzalez Lopez Ledesma, M.M., Ojeda, D.S., Henrion-Lacritick, A., Morales, M.A., Fabri, C., et al. (2020) Zika Virus Subgenomic Flavivirus RNA Generation Requires Cooperativity between Duplicated RNA Structures That Are Essential for Productive Infection in Human Cells. Journal of Virology, 94.

22. Akiyama, B.M., Graham, M.E., O ′ Donoghue, Z. and Jeffrey. (2021) Three-dimensional structure of a flavivirus dumbbell RNA reveals molecular details of an RNA regulator of replication. Nucleic Acids Research, 49, 7122–7138.

23. Graham, M.E., Merrick, C., Akiyama, B.M., Szucs, M.J., Leach, S., Kieft, J.S. and Beckham, J.D. (2023) Zika virus dumbbell-1 structure is critical for sfRNA presence and cytopathic effect during infection. mBio.

24. Liu, Y., Guan, W. and Liu, H. (2023) Subgenomic Flaviviral RNAs of Dengue Viruses. Viruses, 15, 2306.

25. Ward, A.M., Bidet, K., Yinglin, A., Ler, S.G., Hogue, K., Blackstock, W., Gunaratne, J. and Garcia-Blanco, M.A. (2011) Quantitative mass spectrometry of DENV-2 RNA-interacting proteins reveals that the DEAD-box RNA helicase DDX6 binds the DB1 and DB2 3’ UTR structures. RNA Biology, 8, 1173–1186.

26. Ramos-Lorente, S.E., Berzal-Herranz, B., Romero-Lopez, C. and Berzal-Herranz, A. (2024) Recruitment of the 40S ribosomal subunit by the West Nile virus 3’ UTR promotes the cross-talk between the viral genomic ends for translation regulation. Virus Res, 343, 199340.

27. Manzano, M., Reichert, E.D., Polo, S., Falgout, B., Kasprzak, W., Shapiro, B.A. and Padmanabhan, R. (2011) Identification of Cis-Acting Elements in the 3′-Untranslated Region of the Dengue Virus Type 2 RNA That Modulate Translation and Replication. Journal of Biological Chemistry, 286, 22521–22534.

28. Brierley, I., Pennell, S., Gilbert, R. J. (2007) Viral RNA pseudoknots: versatile motifs in gene expression and replication. Nature Reviews Microbiology, 5, 598–610.

29. Wan, H., Jang, H., Xu, L., Abdallah, K.S., Gilbert, W.V. and Pyle, A.M. (2026) A ribosome-bound pseudoknot in the HCV coding region stimulates viral growth by tuning viral translation. Cell Reports, 45, 116739.

30. Chapman, E.G., Moon, S.L., Wilusz, J. and Kieft, J.S. (2014) RNA structures that resist degradation by Xrn1 produce a pathogenic Dengue virus RNA. Elife, 3, e01892.

31. Chapman, E.G., Costantino, D. A., Rabe, J. L., Moon, S. L., Wilusz, J., Nix, J. C., Kieft, J. S.. (2014) The structural basis of pathogenic subgenomic flavivirus RNA (sfRNA) production. Sicence, 344, 307–310.

32. Akiyama, B.M.L., H. M.; Massey, A. R.; Costantino, D. A.; Xie, X.; Yang, Y.; Shi, P.; Nix, J. C.; Beckham, J. D.; Kieft, J. S. (2016) Zika virus produces noncoding RNAs using a multi-pseudoknot structure that confounds a cellular exonuclease. Science, 354, 1148–1152.

33. Guth-Metzler, R., Mohamed, A.M., Cowan, E.T., Henning, A., Ito, C., Frenkel-Pinter, M., Wartell, R.M., Glass, J.B. and Williams, L.D. (2023) Goldilocks and RNA: where Mg2+ concentration is just right. Nucleic Acids Res, 51, 3529–3539.

34. Grilley, D., Misra, V., Caliskan, G., Draper, D. E. (2007) Importance of Partially Unfolded Conformations for Mg2+-Induced Folding of RNA Tertiary Structure: Structural Models and Free Energies of Mg2+ Interactions. Biochemistry, 46.

35. Leipply, D. and Draper, D.E. (2011) Evidence for a Thermodynamically Distinct Mg^2+^ Ion Associated with Formation of an RNA Tertiary Structure. Journal of the American Chemical Society, 133, 13397–13405.

36. Mainan, A., Kundu, R., Singh, R.K. and Roy, S. (2024) Magnesium Regulates RNA Ring Dynamics and Folding in Subgenomic Flaviviral RNA. The Journal of Physical Chemistry B, 128, 9680–9691.

37. Niu, X., Sun, R., Chen, Z., Yao, Y., Zuo, X., Chen, C. and Fang, X. (2021) Pseudoknot length modulates the folding, conformational dynamics, and robustness of Xrn1 resistance of flaviviral xrRNAs. Nature Communications, 12.

38. Liu, T.P., S.; Pyle, A. M. (2023) Making RNA: Using T7 RNA polymerase to produce high yields of RNA from DNA templates. Methods in Enzymology, 691.

39. Tang, G.Q.N., D.; Bandwar, R. P.; Lee, K. S.; Roy, R.; Ha, T.; Patel, S. S. (2014) Relaxed Rotational and Scrunching Changes in P266L Mutant of T7 RNA Polymerase Reduce Short Abortive RNAs while Delaying Transition into Elongation. PLOS ONE, 9.

40. Delgado-Garcia, L., Ambre, S., Filbin, M. and Zimmerman, Z. (2024) Abstract 2085 Optimizing the Procedure for Synthesis of SHAPE Probe 2-methylnicotinic acid imidazolide (NAI). Journal of Biological Chemistry, 300, 106597.

41. Busan, S.W., K. M. (2018) Accurate detection of chemical modifications in RNA by mutational profiling (MaP) with ShapeMapper 2. RNA, 23.

42. Smola, M.J.R., M. G.; Busan, S.; Siegfried, N. A.; Weeks, K. M. (2015) Selective 2′-hydroxyl acylation analyzed by primer extension and mutational profiling (SHAPE-MaP) for direct, versatile and accurate RNA structure analysis. Nature Protocols, 10.

43. Markoff, L. (2003) 5’ and 3’ noncoding regions in Flavivirus RNA.

44. Bellaousov, S., Reuter, J.S., Seetin, M.G. and Mathews, D.H. (2013) RNAstructure: Web servers for RNA secondary structure prediction and analysis. Nucleic Acids Res, 41, W471–474.

45. Johnson, P.Z., Kasprzak, W.K., Shapiro, B.A. and Simon, A.E. (2019) RNA2Drawer: geometrically strict drawing of nucleic acid structures with graphical structure editing and highlighting of complementary subsequences. RNA Biology, 16, 1667–1671.

46. Patel, S., Sexton, A.N., Strine, M.S., Wilen, C.B., Simon, M.D. and Pyle, A.M. (2023) Systematic detection of tertiary structural modules in large RNAs and RNP interfaces by Tb-seq. Nat Commun, 14, 3426.

47. Lanciotti, R.S., Kerst, A. J., Nasci, R. S., Godsey, M. S., Mitchell, C. J., Savage, H. M., Komar, N., Panella, N. A., Allen, B. C., Volpe, K. E., Davis, B. S., Roehrig, J. T. (2000) Rapid detection of west nile virus from human clinical specimens field-collected mosquitoes, and avian samples by a TaqMan reverse transcriptase-PCR assay. Journal of Clinical Microbiology, 38, 4066–4071.

48. Misra, V.K.D., D. E. (2002) The Linkage between Magnesium Binding and RNA Folding. Journal of Molecular Biology, 317.

49. Butcher, S.E. and Pyle, A.M. (2011) The molecular interactions that stabilize RNA tertiary structure: RNA motifs, patterns, and networks. Acc Chem Res, 44, 1302–1311.

50. Soto, A.M., Misra, V. and Draper, D.E. (2007) Tertiary structure of an RNA pseudoknot is stabilized by “diffuse” Mg2+ ions. Biochemistry, 46, 2973–2983.

51. Slonchak, A., Parry, R., Pullinger, B., Sng, J.D.J., Wang, X., Buck, T.F., Torres, F.J., Harrison, J.J., Colmant, A.M.G., Hobson-Peters, J. et al. (2022) Structural analysis of 3’UTRs in insect flaviviruses reveals novel determinants of sfRNA biogenesis and provides new insights into flavivirus evolution. Nature Communications, 13.

52. Khare, B. and Kuhn, R.J. (2022) The Japanese Encephalitis Antigenic Complex Viruses: From Structure to Immunity. Viruses, 14, 2213.

53. Wan, H., Adams, R. L., Lindenbach, B. D., Pyle, A. M. (2022) The *in vivo* and *in vitro* rchitecture of the hepatitis C virus RNA genome uncovers functional RNA secondary and tertiary structures. Journal of Virology, 96.

54. Kieft, J.S., Rabe, J.L. and Chapman, E.G. (2015) New hypotheses derived from the structure of a flaviviral Xrn1-resistant RNA: Conservation, folding, and host adaptation. RNA Biology, 12, 1169–1177.

55. Zhaguparov, D., Zhao, M., Sekar, R.V. and Woodside, M.T. (2025) Identifying the interactions conferring functional mechanical rigidity on RNase-resistant RNA from Zika virus. Proceedings of the National Academy of Sciences, 122.

56. Villordo, S.M., Carballeda, J.M., Filomatori, C.V. and Gamarnik, A.V. (2016) RNA Structure Duplications and Flavivirus Host Adaptation. Trends Microbiol, 24, 270–283.

57. Lai, M.M.C. (2000) The making of infectious viral RNA: No size limit in sight. Proceedings of the National Academy of Sciences, 97, 5025–5027.

58. Baker, C. and Shi, P.Y. (2020) Construction of Stable Reporter Flaviviruses and Their Applications. Viruses, 12.

59. Aubry, F., Nougairede, A., Gould, E.A. and de Lamballerie, X. (2015) Flavivirus reverse genetic systems, construction techniques and applications: a historical perspective. Antiviral Res, 114, 67–85.

60. SERVICES, D.O.H.A.H. (2024) NIH GUIDELINES FOR RESEARCH INVOLVING RECOMBINANT OR SYNTHETIC NUCLEIC ACID MOLECULES (NIH GUIDELINES)

61. Mazeaud, C., Freppel, W. and Chatel-Chaix, L. (2018) The Multiples Fates of the Flavivirus RNA Genome During Pathogenesis. Frontiers in Genetics, 9.

62. Nelson, A.N. and Ploss, A. (2024) Emerging mosquito-borne flaviviruses. mBio, 15.

63. Qassem, S., Breier, D., Naidu, G.S., Hazan-Halevy, I. and Peer, D. (2024) Unlocking the therapeutic potential of locked nucleic acids through lipid nanoparticle delivery. Mol Ther Nucleic Acids, 35, 102224.

64. Huston, N.C., Wan, H., Strine, M. S., de Cesaris Araujo Tavares, R., Wilen, C. B., Pyle, A. M. (2021) Comprehensive in vivo secondary structure of the SARS-CoV-2 genome reveals novel regulatory motifs and mechanisms. Molecular Cell, 81, 584–598.

65. Dethoff, E.A., Boerneke, M. A., Gokhale, N. S., Muhire, B. M., Martin, D. P., Sacco, M. T., McFadden, M. J., Weinstein, J. B., Messer, W. B., Horner, S. M., Weeks, K. M. (2018) Pervasive tertiary structure in the dengue virus RNA genome. Proceedings of the National Academy of the Sciences, 115, 11513–11518.

66. Tuplin, A., Struthers, M., Cook, J., Bentley, K., Evans, D. J. (2015) Inhibition of HCV translation by disrupting the structure and interactions of the viral CRE and 3’ X-tail. Nucleic Acids Research, 43, 2914–2926.

