## Supplementary for "Interdependent RNA structural motifs at the 3ʹ-terminus of the West Nile virus genome regulate viral growth"

Lucille H Tsao *et al.*

*Anna Marie Pyle.

**This PDF file includes:**

Figs. S1 to S7
Tables S1 to S4

### Supplementary Text

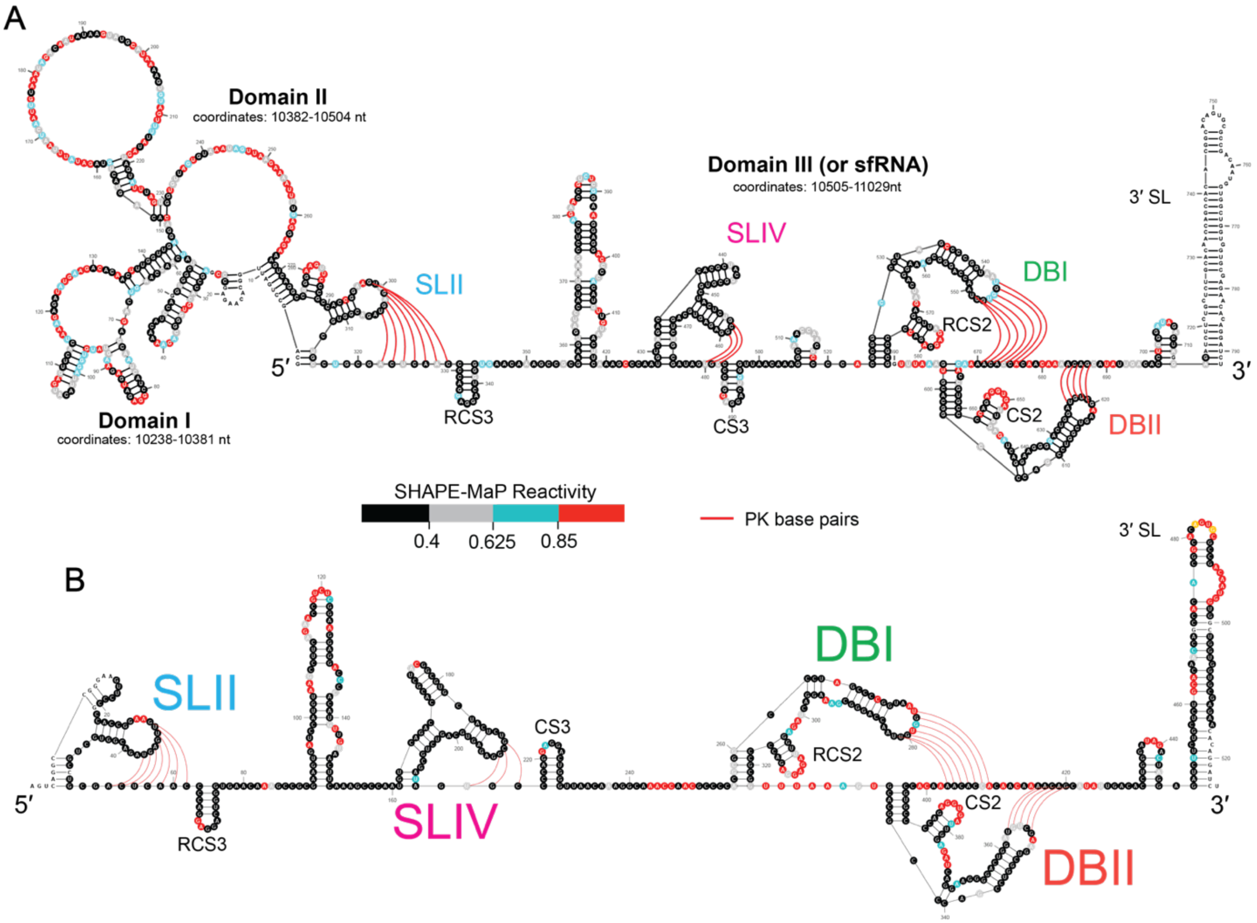


**Fig. S1. Compact secondary structure *in vitro* WNV sfRNA in isolation and WNV 3' viral terminus.** (**A**) The predicted structure of the 3' viral terminus *in vitro*. Nucleotides are colored by normalized SHAPE-MaP reactivities. SHAPE reactivities were categorized based on the following thresholds: <0.40 (black, low levels of modification), 0.40≤ to <0.625 (gray, partial to low levels of modification), 0.625≤ to <0.85 (blue, partial to high levels of modification), ≥0.85 (red, high levels of modification). Pseudoknot base pairs are represented with red lines. (**B**) The average SHAPE-MaP reactivities mapped onto previously published *in silico* and genetically determined model of the sfRNA. Nucleotides are colored by normalized SHAPE-MaP reactivities. SHAPE reactivities were categorized based on the same thresholds as described above. Pseudoknot base pairs are represented with red lines.

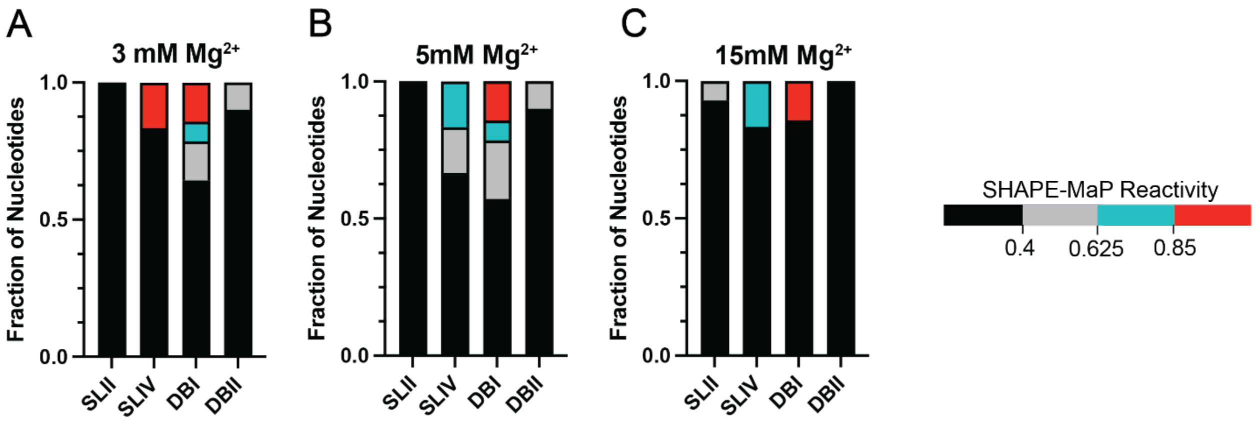


**Figure S2. Pseudoknots form upon addition of $Mg^{2+}$.** (**A-C**) Normalized reactivities for each pseudoknotted nucleotide found in SLII, SLIV, DBI, and DBII PKs were averaged from two replicates at 3, 5, and 15 mM $Mg^{2+}$. Nucleotide reactivities in each PK were then grouped by reactivity level (i.e. amount of SHAPE modification) based on the following thresholds: <0.40 (black, low levels of modification), 0.40≤ to <0.625 (gray, partial to low levels of modification), 0.625≤ to <0.85 (blue, partial to high levels of modification), ≥0.85 (red, high levels of modification). For each PK, the pseudoknotted nucleotides were then expressed in fraction of total plots to display the distribution of SHAPE reactivities found in each PK.

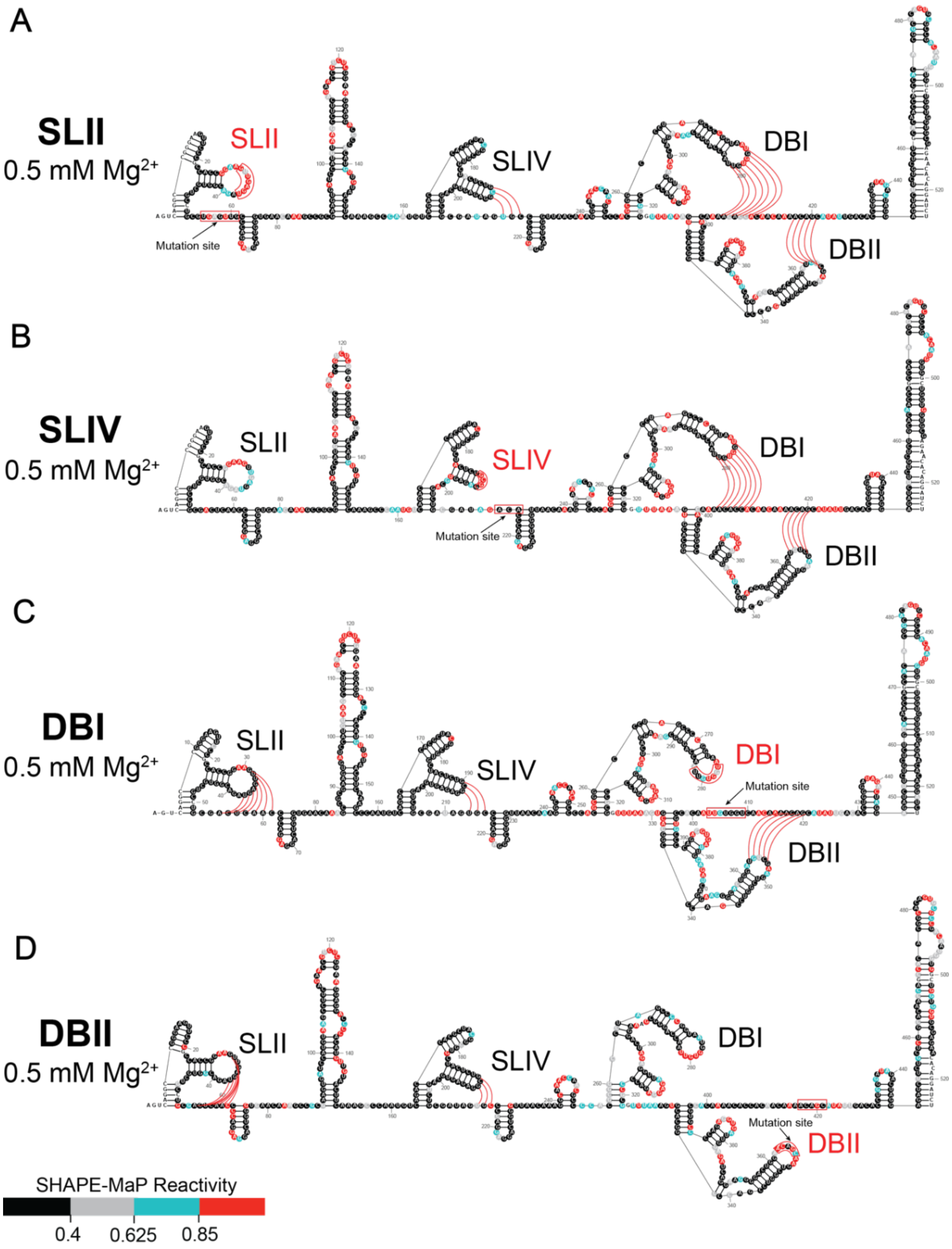


**Figure S3. Superfold prediction of mutant sfRNA transcripts at 0.5 mM $Mg^{2+}$.** (**A-D**) Superfold secondary structure predictions for each sfRNA mutant construct at 0.5 mM $Mg^{2+}$. The unzipped PK in each structure is labeled in red with its PK base-pairing sequences boxed in red. The mutated sequence is labeled as "mutation site." Unchanged PKs are labeled in black, and PK base pairs are indicated with red lines. SHAPE reactivities were categorized based on the following thresholds: <0.40 (black, low levels of modification), 0.40≤ to <0.625 (gray, partial to low levels of modification), 0.625≤ to <0.85 (blue, partial to high levels of modification), ≥0.85 (red, high levels of modification).

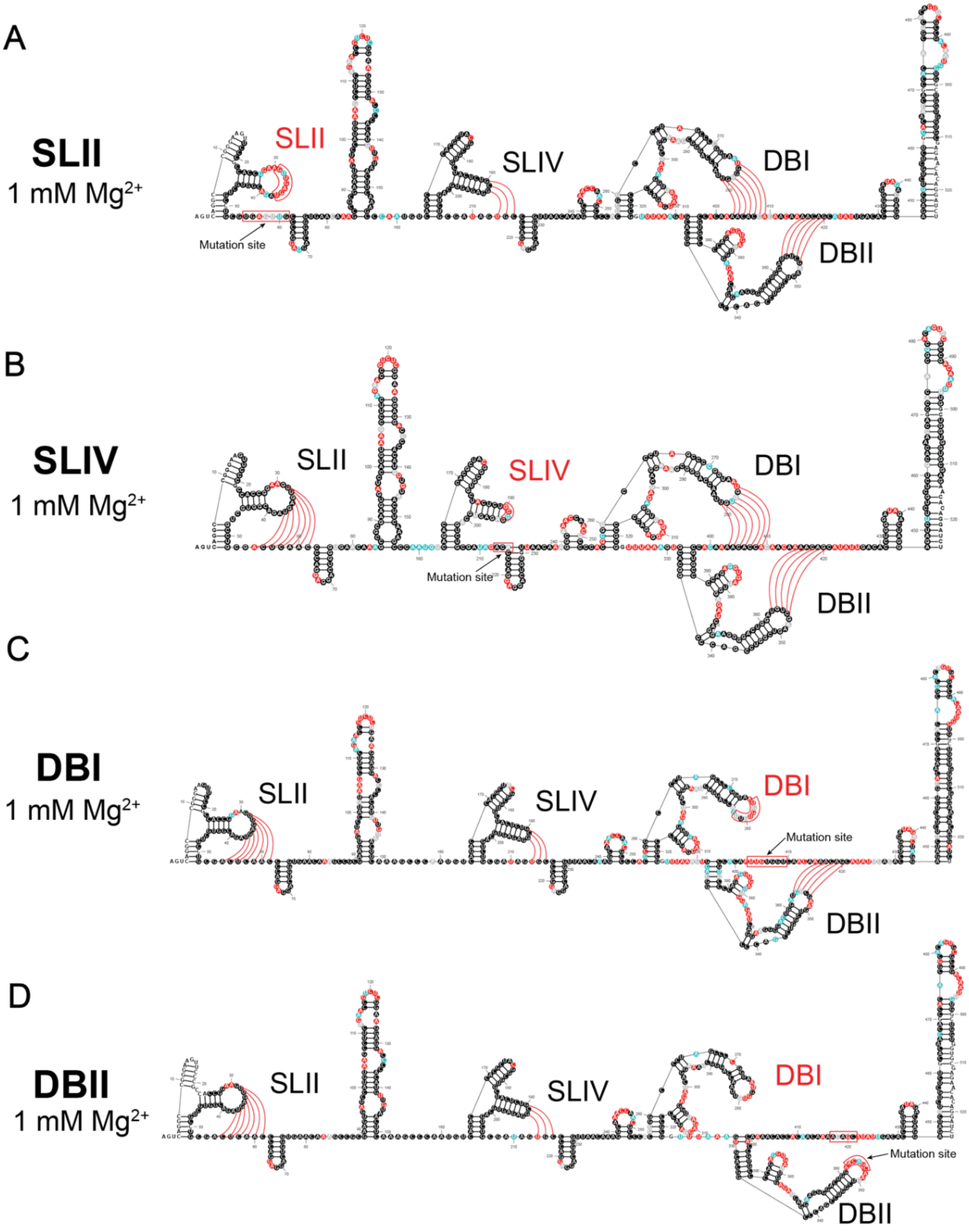


**Figure S4. Superfold prediction of mutant sfRNA transcripts at 1 mM $Mg^{2+}$.** (**A-D**) Superfold secondary structure predictions for each sfRNA mutant construct at 1 mM $Mg^{2+}$. The unzipped PK in each structure is labeled in red with its PK base-pairing sequences boxed in red. The mutated sequence is labeled as "mutation site." Unchanged PKs are labeled in black, and PK base pairs are indicated with red lines. SHAPE reactivities were categorized based on the following thresholds: <0.40 (black, low levels of modification), 0.40≤ to <0.625 (gray, partial to low levels of modification), 0.625≤ to <0.85 (blue, partial to high levels of modification), ≥0.85 (red, high levels of modification).

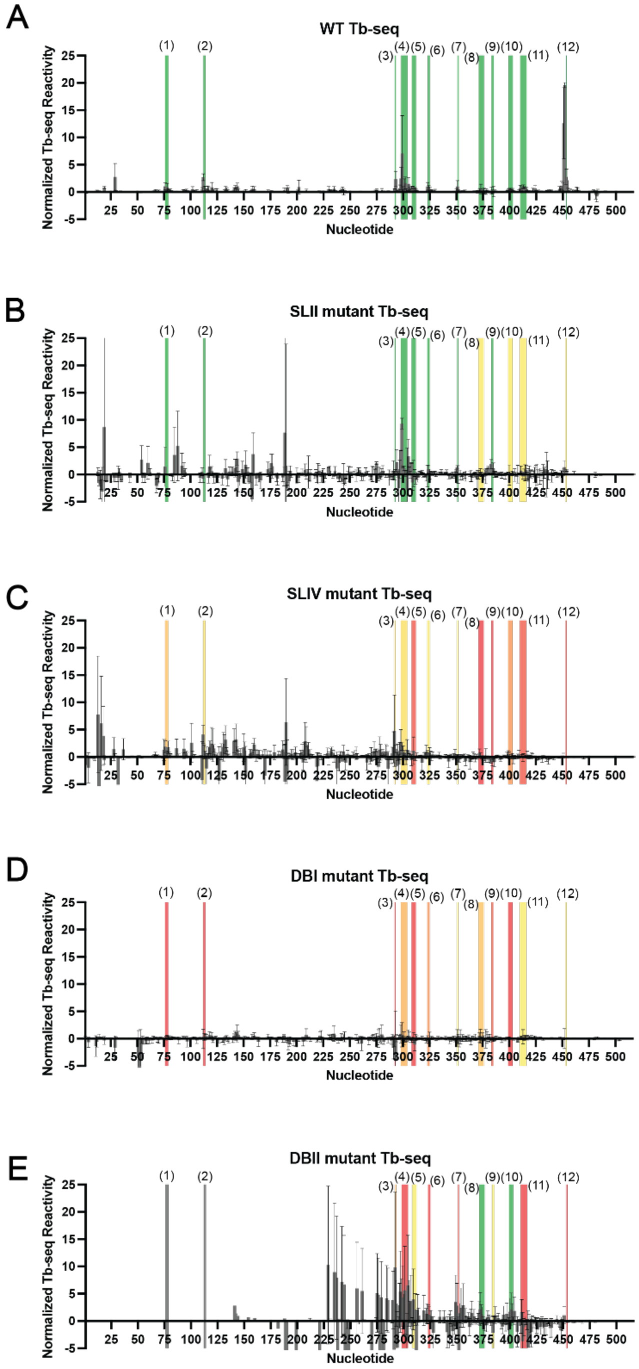


**Figure S5. Locations of tertiary motifs in WT and mutant sfRNAs.** (**A**) Analysis of normalized Tb-seq reactivities (across two replicates) revealed the presence of 12 Tb-sites in the WT sfRNA at 5 mM $Mg^{2+}$. Nucleotide coordinates are displayed on the x-axis. The 12 Tb-sites identified are shaded in green and labeled (1) to (12) in black. (**B-E**) Normalized Tb-seq reactivities of mutant sfRNA constructs at 5 mM $Mg^{2+}$ (across two replicates) reveal whether mutation to a PK will retain or abolish the 12 Tb-sites. Sites that were retained are shaded in green, weakened in yellow, and abolished in red (corresponds to gradient colors in Figure 4).

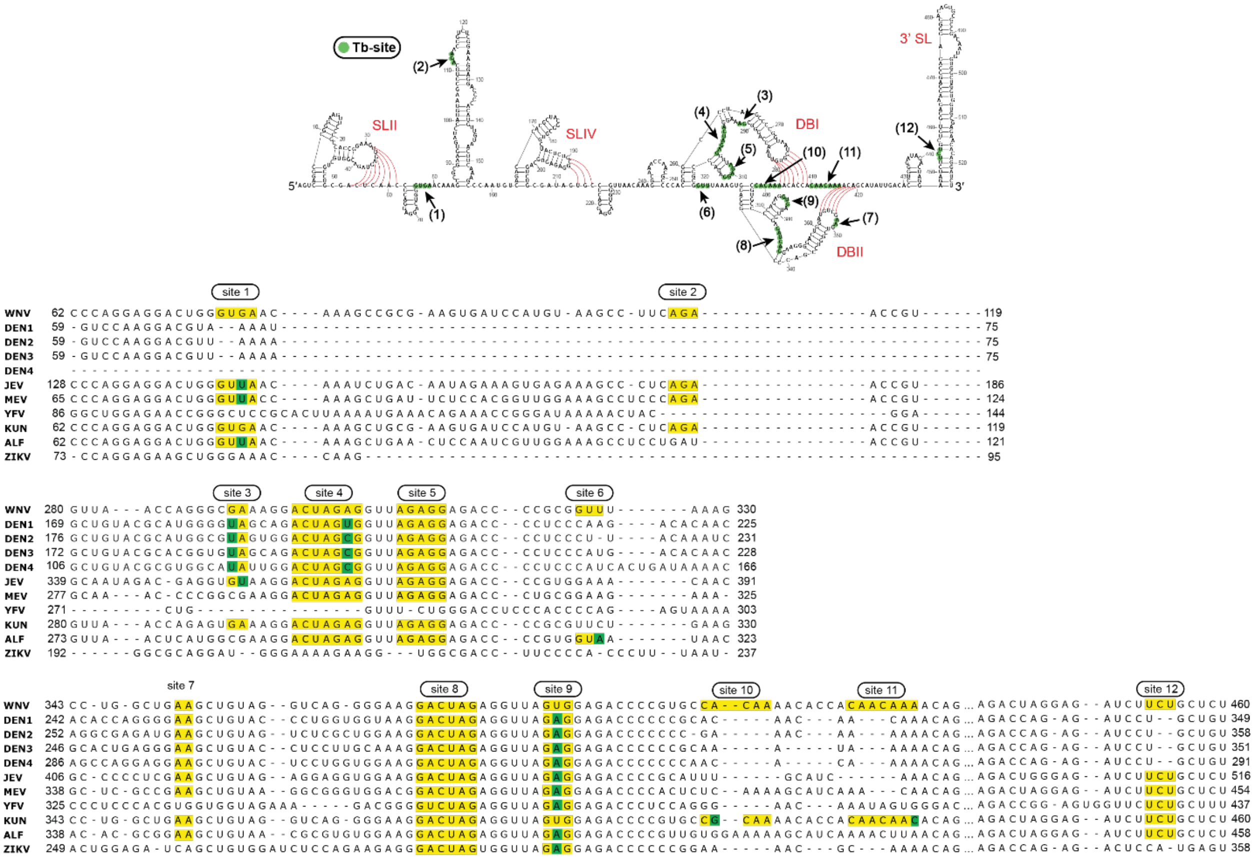


**Figure S6. Tb-sites found WT sfRNA are conserved across multiple flaviviruses.** Schematic showing 12 Tb-sites found in WT sfRNA. Nucleotides found in each Tb-site are highlighted with a green circle. Tb-sites are labeled (1) – (12) in black and indicated with black arrows. PKs are indicated in red, and pseudoknot base pairs are shown with red lines. The sequence conservation of Tb-sites was performed through a multiple sequence alignment was performed using Unipro UGENE Align Tool (Clustal Omega) with WNV serving as the reference sequence. Tb-sites that were found to be 100% conserved in a flavivirus 3' UTR are highlighted in yellow. Single nucleotide differences are highlighted in green.

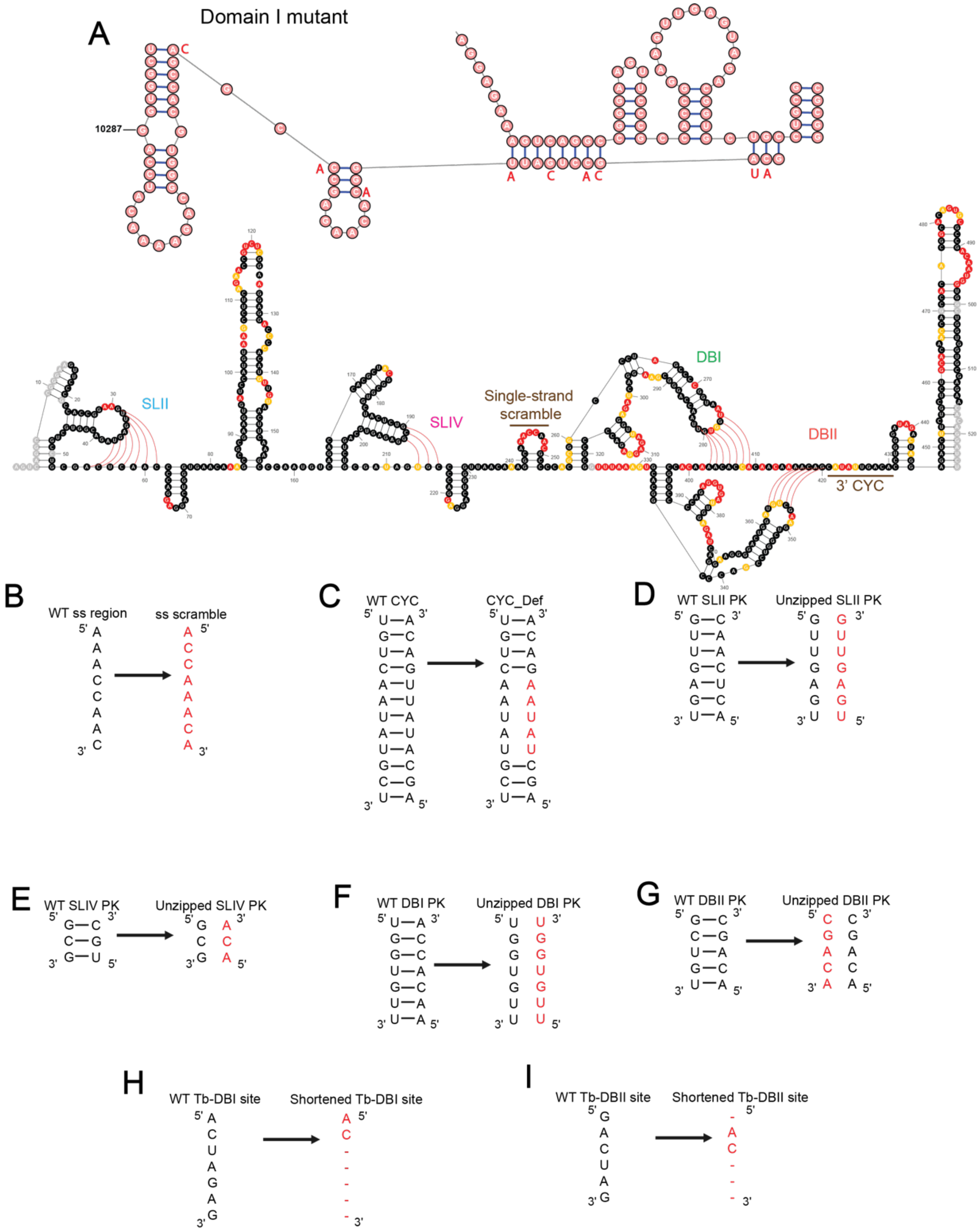


**Figure S7. Mutations to 3' viral terminal structural motifs to assess viral growth.** (**A**) The Domain I mutant was created using synonymous mutations. The specific mutations are noted in red. (**B-I**) Schematic displaying mutations made to each mutant. The right side shows the WT sequence of structural motifs and corresponding base pairs. The left side shows the mutations made to unzip, scramble, or shorten structural motifs. Mutated nucleotides are displayed in red. Deleted nucleotides are noted with "-". More details on mutations found in Methods.

### Table S1. Primers used in this study

| Primer Name | Sequence | Purpose |
|---|---|---|
| **pcDNA3_DIII_T7_R** | GAATTCTGCAGATATCCATCACACTGGCGGCCGC | Cloning |
| **pcDNA3_DIII_EcoRI_R** | CCCTATAGTGAGTCGTATTAATTTCGATAAGC | Cloning |
| **sfRNA_insert_F** | GCTTACTGGCTTATCGAAATTAATACGA | Cloning |
| **sfRNA_insert_R** | GCTCGAGCGGCCGCCAGT | Cloning |
| **sfRNA_509_RT_4** | AGATCCTGTGTTCTCGC | SHAPE-MaP |
| **sfRNA_PCR_F** | AGTCAGGCCGGGAAGTTCC | SHAPE-MaP |
| **STOPSEQ_sfRNA_509_RT_4** | CAGACGTGTGCTCTTCCGATCTAGATCCTGTGTTCTCGC | Tb-seq |
| **3' adaptor** | 5'Phos-NNNNNNAGATCGGAAGAGCGTCGTGTAG-3'Bio | Tb-seq |
| **WNV_1160_F** | TCAGCGATCTCTCCACCAAAG | TaqMan Assay |
| **WNV_1229_R** | GGGTCAGCACGTTTGTCATTG | TaqMan Assay |
| **WNV_1031_F** | TAATACGACTCACTATAGATTTGGTTCTCGAAGGCGACAG | qRT-PCR standard |
| **WNV_3430_R** | GTGGTGGTAAGGTGCAGCTC | qRT-PCR standard |
| **WNV_Env_Taq** | TGCCCGACCATGGGAGAAGCTC | TaqMan Assay probe against ENV, 5'6FAM, 3'quencher = TAMRA |

### Table S2. Summary of quality control metrics from ShapeMapper 2.1.5

| | **Read Depth Check (%)** | **Mutation Rate Check (%)** | **High Background Check (%)** | **Number Highly Reactive (%)** | **Correlation Coefficient (r)** |
|---|---|---|---|---|---|
| **ShapeMapper Thresholds** | **>80%** | **>50%** | **<5%** | **>8%** | |
| WT sfRNA, Rep 1 | 95.4 (501/525) | 80.0 (401/501) | 1.6 (8/501) | 13.4 (67/501) | r = 0.93 |
| WT sfRNA, Rep 2 | 92.8 (487/525) | 70.8 (345/487) | 1.6 (8/487) | 20.9 (102/487) | |
| SLII mutant, Rep 1 | 94.1 (494/525) | 75.9 (375/494) | 1.4 (7/494) | 21.1 (104/494) | r =0.96 |
| SLII mutant, Rep 2 | 92.4 (485/525) | 77.9 (378/485) | 1.6 (8/485) | 27.6 (134/485) | |
| SLIV mutant, Rep 1 | 96.0 (504/525) | 66.7 (336/504) | 1.2 (6/504) | 21.2 (107/504) | r = 0.89 |
| SLIV mutant, Rep 2 | 94.5 (496/525) | 84.1 (417/496) | 1.0 (5/496) | 25.4 (126/496) | |
| DBI mutant, Rep 1 | 97.1 (510/525) | 76.7 (391/510) | 1.4 (7/510) | 9.4 (48/510) | r = 0.94 |
| DBI mutant, Rep 2 | 93.9 (493/525) | 79.7 (393/493) | 1.0 (5/493) | 24.9 (123/493) | |
| DBII mutant, Rep 1 | 91.6 (481/525) | 67.8 (326/481) | 1.2 (6/481) | 15.0 (72/481) | r = 0.83 |
| DBII mutant, Rep 2 | 96.4 (506/525) | 60.3 (305/506) | 1.2 (6/506) | 12.1 (61/506) | |

### Table S3. Constraints used for structure prediction

| Single Strand constraints |
| --- |
| nt coordinates |
| 1 |
| 2 |
| 3 |
| 53 |
| 54 |
| 77 |
| 78 |
| 158 |
| 159 |
| 160 |

| SLII PK constraints | | DBI PK constraints | |
| --- | --- | --- | --- |
| 5' arm, nt coordinates | 3' arm, nt coordinates | 5' arm, nt coordinates | 3' arm, nt coordinates |
| 31 | 61 | 276 | 409 |
| 32 | 60 | 277 | 408 |
| 33 | 59 | 278 | 407 |
| 34 | 58 | 279 | 406 |
| 35 | 57 | 280 | 405 |
| 36 | 56 | 281 | 404 |
| 37 | 55 | 282 | 403 |

| SLIV PK constraints | | DBII PK constraints | |
| --- | --- | --- | --- |
| 5' arm, nt coordinates | 3' arm, nt coordinates | 5' arm, nt coordinates | 3' arm, nt coordinates |
| 190 | 215 | 353 | 421 |
| 191 | 214 | 354 | 420 |
| 192 | 213 | 355 | 419 |
| | | 356 | 418 |
| | | 357 | 417 |

### Table S4. Viral genome sequences used for sequence alignment

| Virus | Abbreviation | Accession Number |
|---|---|---|
| **West Nile virus** | WNV | AF196835.2 |
| **dengue virus 1** | DENV1 | NC_001477.1 |
| **dengue virus 2** | DENV2 | NC_001474.2 |
| **dengue virus 3** | DENV3 | NC_001475.2 |
| **dengue virus 4** | DENV4 | NC_002640.1 |
| **Japanese encephalitis virus** | JEV | AF014161.1 |
| **Murray Valley encephalitis virus** | MEV | NC_000943.1 |
| **yellow fever virus** | YFV | X03700.1 |
| **Kunjin virus** | KUN | L24512.1 |
| **alfuy virus** | ALF | AY898809.1 |
| **Zika virus** | ZIKV | MT636065.1 |
